# A synthetic biology toolkit for interrogating plasmid-dependent methylotrophy and enhancing methanol-based biosynthesis of *Bacillus methanolicus*

**DOI:** 10.1101/2025.05.06.652373

**Authors:** Pan Liu, Qianqian Yuan, Xueting Yang, Qian Wang, Tao Chang, Yaning Bi, Peng Wu, Tong Zhang, Jinxing Yang, Shiting Guo, Chaoyou Xue, Zhaojuan Zheng, Bo Xin, Hongwu Ma, Yu Wang

## Abstract

Bacillus methanolicus represents a thermophilic methylotroph whose methanol utilization depends on plasmid-encoded genes. It serves as a unique model for deciphering plasmid-dependent methylotrophy and an ideal chassis for low-carbon biomanufacturing using CO2-derived C1 substrates. Despite its evolutionary uniqueness and industrial potential, the lack of synthetic biology tools has hindered both mechanistic understanding and strain engineering. Here, we present a comprehensive synthetic biology platform comprising a high-efficiency electroporation protocol, a CRISPR method enabling robust and multiplex genome editing, diverse neutral loci for gene integration and overexpression, and a cloud-based genome-scale metabolic model iBM822 for user-friendly biodesign. Leveraging this toolkit, we systematically dissected plasmid-dependent methylotrophy, host restriction-modification systems, and functional significance of the chromosomal methylotrophic genes through targeted deletion. To address plasmid loss-induced strain degeneration, we integrated the large endogenous plasmid pBM19 into the chromosome for stable and intact methylotrophic growth. Finally, by integrating metabolic modeling with CRISPR editing, we engineered L-arginine feedback regulation to achieve the first L-arginine biosynthesis from methanol. This study establishes a synthetic biology framework for B. methanolicus, promoting mechanistic exploration of methylotrophy and low-carbon biomanufacturing.

## Introduction

Methylotrophy represents the ability to utilize one-carbon (C1) substrates, such as methanol, methane, and other methylated compounds, as the sole carbon and energy sources^1,2^. Since its initial recognition as a metabolic phenomenon in the late nineteenth century, the understanding of methylotrophic metabolism has been continually expanding with the discovery of diverse methylotrophs^3^. Recently, the role of methylotrophs in global biogeochemical processes^4–6^ and low-carbon biomanufacturing has gained significant attention^7–10^. The bioproduction of chemicals, fuels, and foods from CO_2_-hydrogenated C1 feedstocks is considered to be a promising strategy for achieving carbon neutrality^11–14^.

*Bacillus methanolicus* is a plasmid-dependent methylotrophic bacterium, which represents a unique example among diverse methylotrophic species^15^. Most enzymes involved in methylotrophic metabolism are encoded both by an episomal plasmid copy and 1-2 chromosomal copies^16^. The plasmid-dependent methylotrophy is characterized by the fact that iterative cultivation of *B. methanolicus* in methanol-free media leads to the loss of the episomal plasmid, thereby abolishing the ability to grow on methanol^17^. Conversely, the biological role of the chromosomal copies of methylotrophic genes remains unclear. *B. methanolicus* is also a thermophilic and halotolerant bacterium with rapid growth at 50-55°C and in sea water-based media^18^. The application of extremophiles in biomanufacturing is expected to reduce bioreactor cooling requirements, lower contamination risks, increase reaction efficiency, and save freshwater resources^19^. Therefore, *B. methanolicus* is considered as a promising microbial chassis for biomanufacturing using C1 feedstocks^20^. However, the product portfolio of *B. methanolicus* is still limited and its bioproduction levels are significantly lower compared to glucose-based processes using systematically engineered model microorganisms such as *Escherichia coli* and *Saccharomyces cerevisiae*^20–22^.

To elucidate the molecular basis of its unique features and reprogram its metabolism for stable and efficient bioproduction from C1 feedstocks, there is an urgent need for a synthetic biology toolkit for *B. methanolicus*. This toolkit should include CRISPR tools for robust genome editing, neutral loci for gene integration and overexpression, and user-friendly genome-scale metabolic models (GEMs) for biodesign. ^23–25^ However, electroporation of *B. methanolicus* typically exhibits low efficiency^15,17^. Although the preparation of plasmids in *Geobacillus thermoglucosidasius* for methylation can enhance the plasmid transformation efficiency of *B. methanolicus*, these additional steps complicate the procedure^26^. Multi-step recombination with suicide plasmid has historically been used for genetic manipulation^26,27^. However, recent methods still require extremely large homologous arms (HAs, 6 kb) and time-consuming procedures, hindering their application in large-scale and iterative genome editing^26^. Recently, CRISPR has revolutionized the ability to genetically modify microorganisms^28,29^. CRISPR interference using the catalytically deactivated *Streptococcus pyogenes* Cas9 (dSpCas9) has been successfully applied to repress gene expression in *B. methanolicus*^30^, whereas CRISPR-enabled genome editing tools are still unavailable. Additionally, no GEM of *B. methanolicus* has been constructed yet, impeding the design–build–test–learn (DBTL) cycle of synthetic biology.

In this study, we develop a modular synthetic biology toolkit for *B. methanolicus*, and demonstrate its applicability for interrogating gene function, chromosome engineering, and designing and enhancing biosynthetic pathways. Our toolkit includes a simple protocol for highly efficient electroporation, a CRISPR method for robust and multiplex deletion and integration of large-size DNAs, diverse neutral loci for targeted gene integration and overexpression, and a cloud-based GEM for pathway visualization and design. We use these tools to delete episomal plasmids and chromosomal genes to clarify their functions in methylotrophic growth and restriction-modification (RM). Additionally, we integrate the episomal plasmid containing methylotrophic genes into the chromosome for stable and intact methylotrophic growth without degeneration. Furthermore, we develop a GEM and user-friendly cloud-based tools for biodesign, enabling the simulation of biosynthesis from methanol and glucose. Finally, we decipher and release the feedback regulation of L-arginine biosynthesis for L-arginine overproduction from methanol. This work provides synthetic biology tools and approaches that pave the way for systematic engineering of *B. methanolicus* for understanding methylotrophy and promoting methanol-based biomanufacturing.

## Results

### Optimizing the electroporation procedure for efficient DNA transformation

A protocol for DNA transformation via electroporation was developed^15,17^, and a transformation efficiency of 5.3 × 10^2^ cfu/μg DNA was obtained for the *E. coli*-*B. methanolicus* shuttle vector pNW33N (GenBank: AY237122.1) in our test (Fig. 1a). During the preparation of competent cells, we noticed that the EP buffer (1 mM HEPES, 25% polyethylene glycol (PEG) 8000, pH 7.0) for washing and resuspending is very viscous because of PEG8000. Significant cell loss was observed during the washing step. Therefore, PEG8000 was removed from the washing buffer and only used for resuspension of competent cells after the washing step. This modification led to a 10-fold increase in transformation efficiency (5.5 × 10^3^ cfu/μg DNA) (Fig. 1a). PEG with different molecular weights (2000 to 8000) and different concentrations (15% to 35%) did not affect transformation efficiency (Supplementary Fig. 1 and 2).

**Figure 1.**
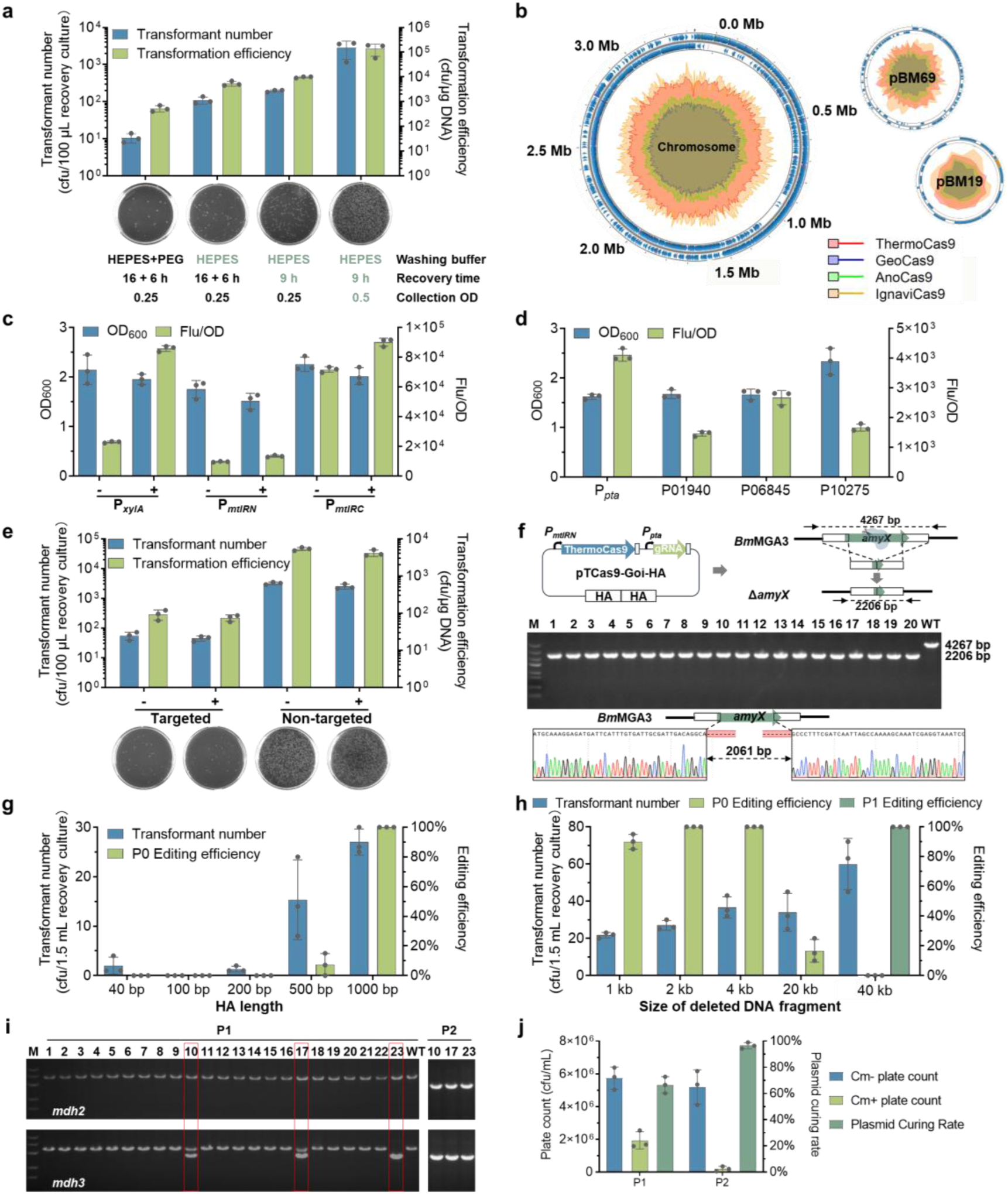
A thermostable CRISPR/Cas9 system for efficient genome editing in *B. methanolicus*. **a**, Systematic optimization of the DNA electroporation procedure. **b,** Comparison of thermostable CRISPR systems based on the number of targetable sites (available PAMs) in the *B. methanolicus* chromosome and episomal plasmid. Coding sequences (CDSs) are shown in blue arrows. The number of targetable sites (available PAMs) in the chromosome and plasmid are counted every 10 kb and 1 kb, respectively, and displayed using radar charts. **c,** Evaluation of inducible promoters for ThermoCas9 expression in *B. methanolicus*. The symbols “+” and “–” indicate the presence or absence of inducers (xylose for P*_xylA_* and mannitol for P*_mtlRN_* and P*_mtlRC_*), respectively. **d,** Evaluation of constitutive promoters for gRNA expression in *B. methanolicus*. **e,** Lethality induced by CRISPR/ThermoCas9-mediated double-strand breaks. The symbols “+” and “–” denote the presence or absence of the mannitol inducer, respectively. “Targeted” and “Non-targeted” refer to plasmids expressing chromosome-targeted and non-targeted gRNAs, respectively. **f,** Deletion of *amyX* gene using CRISPR/ThermoCas9 with 1 kb HAs. **g,** Deletion of *amyX* gene using CRISPR/ThermoCas9 with HAs of varying lengths. **h,** Deletion of DNA fragments of different sizes. P0 represents the editing efficiency directly determined by colony PCR of the transformants of ThermoCas9 plasmid. P1 represents the editing efficiency after one passage cultivation. **i,** Simultaneous deletion of *mdh2* and *mdh3* genes. P0 represents the editing efficiency directly determined by colony PCR of the transformants of ThermoCas9 plasmid. P1 represents the editing efficiency after one passage cultivation. **j,** Curing efficiency of the ThermoCas9-expressing plasmid after one (P1) and two (P2) passages of cultivation in the absence of antibiotics. Values and error bars represent the mean ± standard deviation (s.d.) of three biological replicates (n = 3).

The original procedure required two passages of recovery after electroporation, 16 h and 6 h, respectively. This prolonged cultivation is time-consuming and may result in cells entering the decline phase. We found that only one passage of cultivation for 9 h increased the transformation efficiency to 9.8 × 10^3^ cfu/μg DNA (Supplementary Fig. 3). When preparing competent cells, cells were harvested at a very low optical density at 600 nm (OD_600_) of 0.25. We found that collecting cells at OD_600_ of 0.5 not only increased the competent cell yields per batch but also enhanced the transformation efficiency to 2.2 × 10^5^ cfu/μg DNA (Supplementary Fig. 4). All the transformation tests were performed using 1 μg plasmid DNA. Increasing the usage of DNA to 2 μg or decreasing it to 0.2 μg did not significantly affect the number of transformants, whereas the use of only 0.2 μg led to a higher calculated transformation efficiency of 6.9 × 10^5^ cfu/μg DNA (Supplementary Fig. 5). In summary, with these modifications to the original procedure, the electro-transformation efficiency was increased by approximately 1000-fold (Fig. 1a), laying a foundation for genetic manipulation of *B. methanolicus*.

### Adapting a thermostable CRISPR/Cas9 system for multiplex genome editing

Due to the limited success in adapting CRISPR systems for genome editing in *B. methanolicus*, we designed and developed a CRISPR/Cas9-based approach from the ground up. The initial step involved selecting an appropriate CRISPR/Cas9 system. The widely used SpCas9 is inactive *in vivo* at temperatures above 42°C^31^, and *B. methanolicus* exhibits extremely slow growth under such conditions^32^. Consequently, thermostable CRISPR/Cas9 systems were prioritized. Several such systems derived from thermophiles have been identified, including ThermoCas9 from the *G. thermodenitrificans*^33^, GeoCas9 from *G. stearothermophilus*^34^, IgnaviCas9 from uncultured Ignavibacterium^35^, and AnoCas9 from *Anoxybacillus flavithermus*^36^. These thermostable CRISPR systems exhibit distinct protospacer adjacent motif (PAM) preferences, resulting in varying numbers of targetable sites in the chromosome and plasmids of *B. methanolicus*. ThermoCas9 and IgnaviCas9 target similar number of sites in the chromosome and two endogenous plasmids, 157,302 and 137,576 sites, respectively (Fig. 1b). AnoCas9 and GeoCas9 target significantly fewer sites, with 93,728 and 75,471 targetable sites, respectively. Among these thermostable CRISPR/Cas9 systems, ThermoCas9^33^ and GeoCas9^34^ have been validated for *in vivo* genome editing. Based on the number and distribution of targetable sites, ThermoCas9 was selected for application in *B. methanolicus*.

Subsequently, suitable promoters were identified for the expression of ThermoCas9 and gRNA in *B. methanolicus*. Drawing on experience from developing genome editing tools for recalcitrant non-model microorganisms^37^, inducible and constitutive promoters were employed for the expression of ThermoCas9 and gRNA, respectively. Mannitol- and xylose-inducible promoters were evaluated using the superfolder green fluorescent protein (sfGFP) as a reporter in *B. methanolicus*^38,39^. Upon the addition of xylose, the P*_xylA_* promoter exhibited a 3.8-fold induction (Fig. 1c). The mannitol-inducible promoter P*_mtlR_*_N_ displayed very low expression levels and a 1.4-fold induction. Substitution of the native ribosome-binding site (RBS) of P*_mtlRN_* (AGTGGAG) with the *B. methanolicus* consensus RBS motif (AGGAGG) generated a recombinant P*_mtlRC_* promoter. The expression level increased significantly by 6.8-fold, while only a 1.3-fold induction was observed with mannitol (Fig. 1c). During the construction of the ThermoCas9 expression plasmid in *E. coli*, only the P*_mtlR_*_N_-ThermoCas9 plasmid was successfully generated, while the other two constructs failed. This outcome may be attributed to the high-copy-number ColE1 origin of pNW33N in *E. coli* and the elevated basal expression levels of P*_xylA_* and P*_mtlRC_* compared to P*_mtlRN_* (Supplementary Fig. 6). Three native strong constitutive promoters from *B. methanolicus* (BMMGA3_01940, BMMGA3_06845, and BMMGA3_10275) were evaluated for gRNA expression^40^. However, these native promoters exhibited lower activity compared to the constitutive *pta* promoter (P*_pta_*) from *B. coagulans* in *B. methanolicus*^33^ (Fig. 1d). Based on these findings, P*_mtlRN_* and P*_pta_* were selected to regulate the expression of ThermoCas9 and gRNA, respectively.

To assess the nuclease activity of ThermoCas9 in introducing double-stranded DNA breaks in *B. methanolicus*, a chromosomal gene-targeting gRNA and a non-targeting control gRNA were co-expressed with ThermoCas9 on pNW33N. Chromosome-targeting cleavage resulted in a lethality rate of 98% (Fig. 1e). The addition of mannitol did not enhance lethality, consistent with the similar sfGFP expression levels observed with or without mannitol induction (Fig. 1c). These findings demonstrate the functional expression of ThermoCas9 and gRNA in *B. methanolicus*.

By integrating two 1 kb HAs into the ThermoCas9 and gRNA-expressing plasmid, the deletion of the ∼2 kb *amyX* gene (GeneID: BMMGA3_13410, encoding pullulanase) was evaluated. Colony PCR analysis of 20 transformants revealed that all transformants (Passage 0) exhibited successful editing with a 100% success rate, eliminating the need for further cultivation (Fig. 1f). This streamlined the genome editing procedure, making it both simple and efficient. Subsequently, shorter HAs ranging from 40 bp to 500 bp were tested. Gene deletion was achieved only with 500 bp HAs, albeit with an efficiency of less than 10% (Fig. 1g). Consequently, 1 kb HAs were employed in subsequent experiments. The developed ThermoCas9 system was further utilized to delete DNA fragments of varying sizes. Deletion of DNA fragments up to ∼4 kb in size was achieved with editing efficiencies exceeding 90% (Fig. 1h). However, when targeting a ∼20 kb DNA fragment, the editing efficiency dropped below 10%. For the deletion of a ∼40 kb DNA fragment, no editing events were detected in the transformants (Passage 0). To address this, the transformants were suspended in 0.85% NaCl solution and plated for a second round of cultivation (Passage 1). Analysis of 20 newly grown colonies demonstrated 100% deletion efficiency (Fig. 1h), highlighting the capability of the developed CRISPR tool for manipulating large DNA fragments in *B. methanolicus*.

Multiplex genome editing represents a powerful strategy to significantly accelerate strain engineering^41^. To demonstrate this capability, we targeted two methanol dehydrogenase (Mdh)-encoding genes, *mdh2* (GeneID: BMMGA3_03335) and *mdh3* (GeneID: BMMGA3_09375), and provided two pairs of HAs for simultaneous deletion. Initial colony PCR analysis of the transformants (Passage 0) indicated successful editing of *mdh3* in 3 out of 23 transformants, while *mdh2* remained unedited. Following the same approach used for deleting the ∼40 kb DNA fragment, the three *mdh3*-edited transformants were suspended in 0.85% NaCl solution and subjected to a second round of cultivation (Passage 1). Subsequent colony PCR of the newly obtained colonies confirmed 100% successful double deletion of *mdh2* and *mdh3* (Fig. 1i). The ability to manipulate large DNA fragments and perform multiplex genome editing underscores the versatility and efficiency of the adapted CRISPR system, establishing it as a robust tool for studying and engineering *B. methanolicus*.

Following genome editing, the ThermoCas9 plasmid must be cured to generate an engineered strain suitable for downstream applications or subsequent editing rounds. To achieve this, an edited colony was cultured in liquid media without chloramphenicol for serial passages. The cultivation temperature was increased to 58°C to repress plasmid replication. Mannitol was added to induce ThermoCas9 expression, which was expected to impose a metabolic burden and inhibit cell growth. Consequently, cells that lost the ThermoCas9 plasmid gained a growth advantage in the culture. After two and four passages of cultivation, the plasmid curing rates reached 67% and 97%, respectively (Fig. 1j).

### Investigating plasmid-dependent methylotrophy and RM

The *B. methanolicus* type strain MGA3 harbors two endogenous plasmids, pBM19 and pBM69. pBM19 contains 22 coding sequences (CDSs) and 3 non-coding RNAs (ncRNAs), including six methylotrophic genes (*mdh*^P^, *rpe*^P^, *pfk*^P^, *glpX*^P^, *fba*^P^, and *tkt*^P^)^42^. pBM69 comprises 82 CDSs and 2 ncRNAs, including genes encoding a restriction endonuclease *Bme*TI (GeneID: BMMGA3_16960, an isoschizomer of *Bcl*I recognizing the sequence TGATCA) and a DNA methylase (GeneID: BMMGA3_16950)^43^. Previous studies suggest that this RM system may influence DNA transformation efficiency in *B. methanolicus*^42,43^, although this hypothesis has not been experimentally validated. Notably, the physiological functions of most genes on these plasmids remain uncharacterized. The development of a CRISPR-based genome editing tool has enabled the curing of these two episomal plasmids, facilitating the investigation of their roles in methylotrophy, RM, and potentially other biological processes.

Curing of endogenous plasmids does not require homologous recombination but relies solely on gRNA-guided plasmid cleavage. Two *B. methanolicus* mutants, ΔpBM19 and ΔpBM69, were successfully generated and evaluated for cell growth and DNA transformation efficiency (Fig. 2a). In TSB rich medium without methanol, both ΔpBM19 and ΔpBM69 mutants exhibited slightly slower growth compared to the wild-type strain (Fig. 2b). The ΔpBM19 mutant lost the ability to grow on methanol but demonstrated similar growth on mannitol relative to the wild-type strain (Fig. 2b). To complement the methylotrophic genes on pBM19, the plasmid was extracted from *B. methanolicus* and modified by incorporating a low-copy pSC101 origin and a kanamycin resistance gene for replication in *E. coli* (Fig. 2c). The resulting plasmid, pBMEC22, was successfully transformed into the *B. methanolicus* ΔpBM19 mutant. The recombinant strain regained the ability to grow solely on methanol and exhibited slightly faster growth compared to the wild-type strain (Fig. 2b).

**Figure 2.**
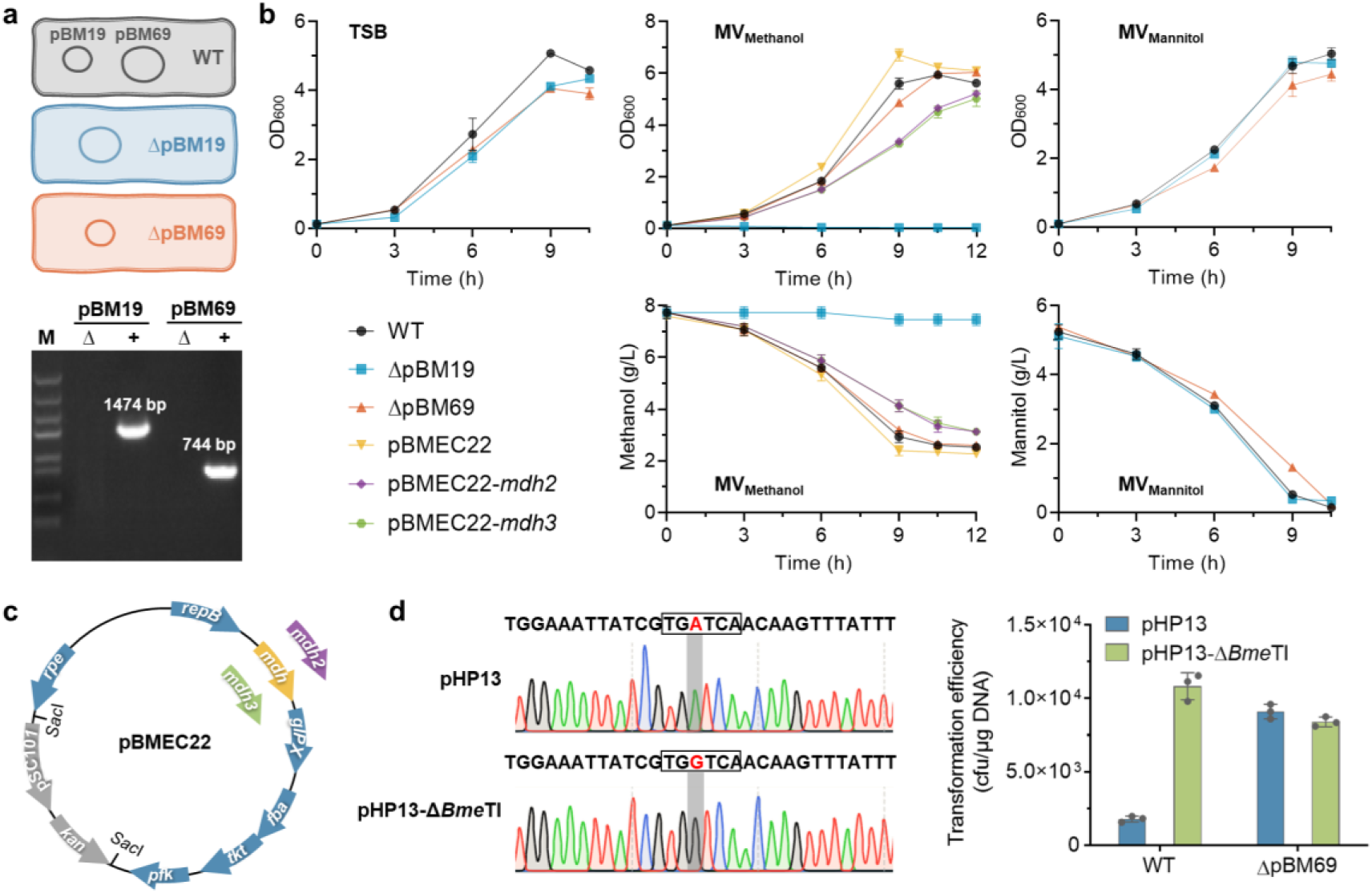
CRISPR-mediated curing and function analysis of endogenous plasmids. **a,** PCR verification of the curing of endogenous plasmids pBM19 and pBM69. **b,** Cell growth of strains engineered with endogenous plasmids in different media. MV_Methanol_ and MV_Mannitol_ represent minimal media containing methanol and mannitol as the sole carbon source, respectively. **c,** Plasmid maps of pBMEC22, pBMEC22-*mdh2*, and pBMEC22-*mdh3*, derivatives of pBM19 capable of replication in both *E. coli* and *B. methanolicus*. **d,** Effects of the pBM69-encoded RM system on the transformation of DNA lacking methylation. The pHP13 plasmid, containing a single *Bme*TI recognition site, and its derivative pHP13-Δ*Bme*TI, with the *Bme*TI recognition site deleted, were prepared in *E. coli* Trans110 (*dam*^−^) and used for transformation experiments. Values and error bars represent the mean ± standard deviation (s.d.) of three biological replicates (n = 3).

The pBMEC22 plasmid can be systematically modified through the deletion or replacement of pBM19-borne genes, either individually or in combination. The *B. methanolicus* ΔpBM19 mutant strain, along with the pBMEC22 plasmid, serves as a valuable tool for investigating gene function. To demonstrate this utility, we replaced the *mdh*^P^ gene in pBMEC22 with the chromosomal *mdh2* and *mdh3* genes, respectively. The resulting plasmids, pBMEC22-*mdh2* and pBMEC22-*mdh3*, successfully restored methylotrophic growth in the *B. methanolicus* ΔpBM19 mutant, whereas the growth was slightly slower than the wild-type strain (Fig. 2b). This finding suggests that *mdh*^P^, *mdh2*, and *mdh3* share similar physiological function in methanol oxidation when expressed at comparable levels.

To investigate the function of the RM system encoded by pBM69, we constructed a new plasmid lacking the *Bme*TI recognition site (TG<u>A</u>TCA mutated to TG<u>G</u>TCA) based on the *E. coli-B. methanolicus* shuttle vector pHP13, which contains a single *Bme*TI recognition site (GenBank: DQ297764.1). Both pHP13 and pHP13-Δ*Bme*TI plasmids were transformed into *E. coli* Trans110 (*dam*^−^) to generate plasmids devoid of m^6^A methylation. The transformation efficiency of pHP13-Δ*Bme*TI into the *B. methanolicus* wild-type strain was 6-fold higher than that of pHP13 (Fig. 2d), indicating that the pBM69-encoded RM system functions as a defense mechanism against DNA lacking m^6^A methylation. When pBM69 was cured, no significant difference in transformation efficiency was observed between pHP13 and pHP13-Δ*Bme*TI (Fig. 2d). Given that pBM69 encodes a functional RM system and its curing does not significantly impact cell growth across all tested media, the *B. methanolicus* ΔpBM69 strain represents a promising new chassis for bioengineering applications.

### Interrogating the function of chromosomal methylotrophic genes

In *B. methanolicus*, methylotrophic gene copies are present on both the episomal plasmid pBM19 and the chromosome (Fig. 3a). While the pBM19 plasmid is considered essential for methanol metabolism, the functional significance of the chromosomal methylotrophic genes remains unclear. To address this, we utilized the developed CRISPR system to individually delete 10 chromosomal genes potentially involved in methylotrophy: *mdh2*, *mdh3*, *act*, *rpe*, *hps*, *phi*, *pfk*, *glpX*, *fba*, and *tkt*. All the ten genes were successfully deleted with high efficiencies ranging from 44% to 100% (Fig. 3b), suggesting the robustness of the developed CRISPR editing tool. Additionally, *mdh2* and *mdh3*, both encoding Mdh, were simultaneously deleted using multiplex genome editing.

**Figure 3.**
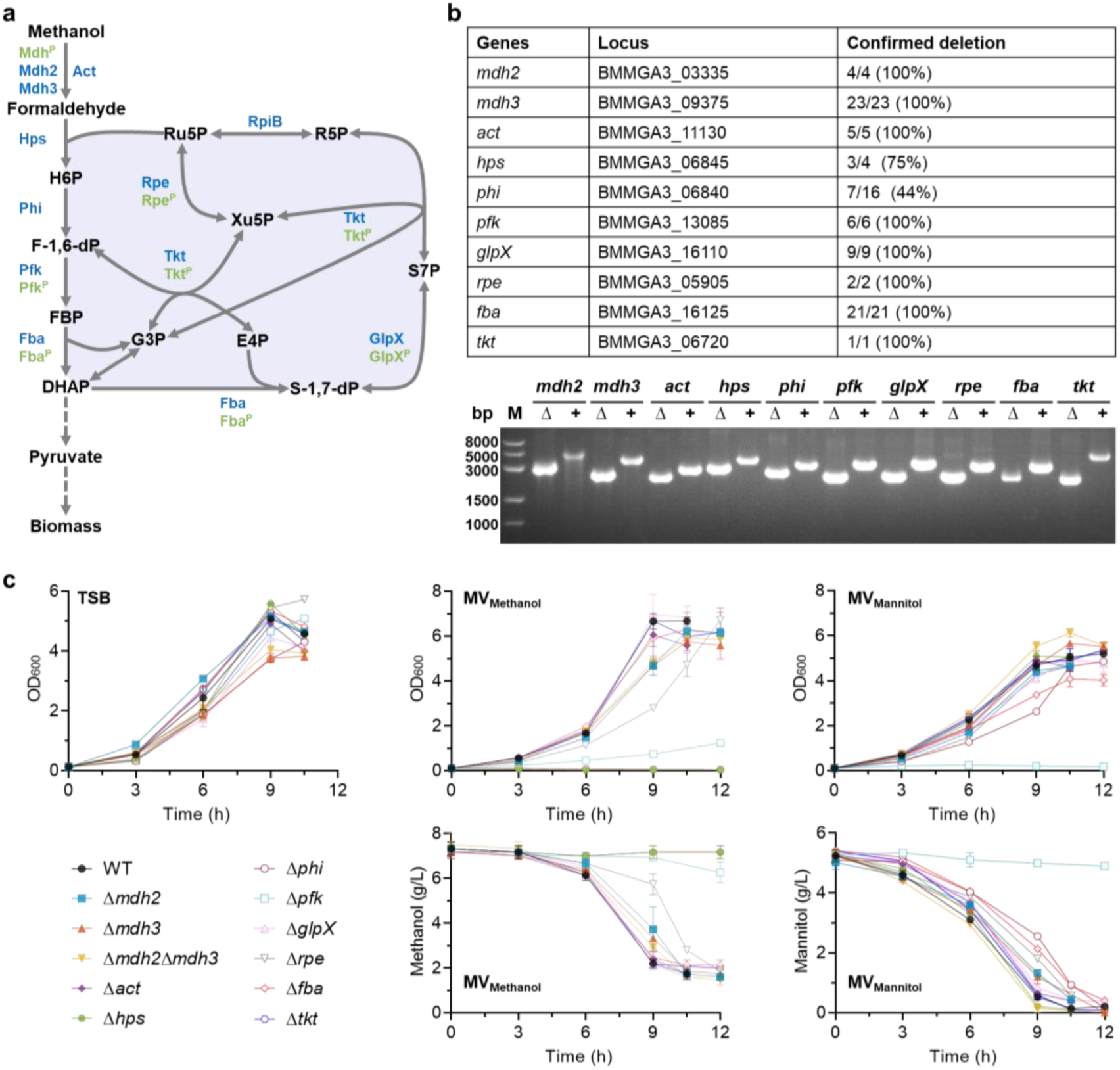
Deletion and function analysis of chromosomal methylotrophic genes. **a,** Schematic representation of the methanol assimilation pathway and the multicopy nature of methylotrophic genes. Enzyme: Act, activator protein of Mdh; Mdh, Mdh2, and Mdh3, methanol dehydrogenase; Hps, 3-hexulose-6-phosphate synthase; Phi, 6-phospho-3-hexuloisomerase; Pfk, 6-phosphofructokinase; Fba, fructose-bisphosphate aldolase/sedoheptulose-bisphosphate aldolase; Tkt, transketolase; Rpe, ribulose-phosphate 3-epimerase; RpiB, ribose-5-phosphate isomerase; GlpX, bifunctional fructose bisphosphatase/sedoheptulose bisphosphatase. Enzymes marked with P are encoded by genes in the episomal plasmid pBM19. Metabolites: Ru5P, ribulose-5-phosphate; R5P, ribose-5-phosphate; Xu5P, xylulose-5-phosphate; H6P, hexulose-6-phosphate; F6P, fructose-6-phosphate; F-1,6-dP, fructose-1,6-bisphosphate; DHAP, dihydroxyacetone phosphate; G3P, glyceraldehyde-3-phosphate; E4P, erythrose-4-phosphate; S7P, sedoheptulose-7-phosphate; S-1,7-dP, sedoheptulose-1,7-bisphosphate. **b,** Deletion of chromosomal methylotrophic genes. Δ and + represent gene-deleted mutant and wild-type strain, respectively. **c,** Cell growth of gene-deleted strains in different media. MV_Methanol_ and MV_Mannitol_ represent minimal media containing methanol and mannitol as the sole carbon source, respectively. Values and error bars represent the mean ± standard deviation (s.d.) of three biological replicates (n = 3).

In TSB rich medium without methanol, most of the gene-deleted mutants exhibited growth rates comparable to the wild-type strain, except that the Δ*mdh3*, Δ*mdh2*Δ*mdh3*, and Δ*phi* mutants showed slightly slower growth (Fig. 3c). More pronounced differences in cell growth were observed in methanol minimal medium. Both individual and combinatorial deletions of *mdh2* and *mdh3* resulted in a slight reduction in cell growth. Previous studies have proposed that Mdh2 and Mdh3 may primarily function in the oxidation of alternative carbon substrates rather than methanol^44^. Our findings suggest that these enzymes also contribute to methanol metabolism, albeit not as the dominant Mdh for methanol oxidation. Interestingly, although Mdh^P^, Mdh2, and Mdh3 are all catalytically enhanced by the activator protein Act, with activity increases of up to 10-fold^45^, the deletion of *act* had no significant impact on cell growth in methanol medium (Fig. 3c). This indicates that Act plays a non-essential role *in vivo*.

Deletion of the chromosomal *rpe* gene significantly impaired cell growth and reduced methanol uptake, while disruption of *pfk* nearly abolished methanol metabolism. Biochemical assays of heterologously expressed enzymes confirmed that both the chromosome- and plasmid-encoded Rpe and Pfk are functional^46^. These results demonstrate that the chromosomal copies of *rpe* and *pfk* are critical for methylotrophy. In contrast, deletion of the chromosomal *glpX*, *fba*, and *tkt* did not affect methanol metabolism, suggesting that the corresponding plasmid-encoded enzymes are sole physiologically functional for methylotrophy. Deletion of either *hps* or *phi* completely abolished methanol metabolism (Fig. 3c), consistent with their exclusive presence in the chromosome. We further evaluated the effects of these gene deletions on growth and metabolism in mannitol medium. Disruption of *fba* and *phi* moderately inhibited cell growth in mannitol minimal medium, whereas *pfk* deletion completely abolished mannitol metabolism (Fig. 3c). These findings indicate that the chromosomal Pfk is essential for both methanol and mannitol metabolism.

### Identification and evaluation of neutral loci for gene integration and expression

Previous metabolic engineering efforts in *B. methanolicus* have predominantly relied on plasmid-based gene expression systems^20^. However, such systems are unsuitable for large-scale and long-term production processes due to their dependence on antibiotic selection markers and inherent segregational instability^47^. Furthermore, microbial heterogeneity arising from plasmid loss and mutation often leads to the emergence of low- and non-productive cell subpopulations during biosynthesis^48^. The developed CRISPR-based tools have enabled the integration of heterologous genes or gene clusters into the *B. methanolicus* chromosome, thereby facilitating stable gene expression. A critical challenge in this regard is the identification of appropriate genomic loci for gene integration. Neutral loci, which permit targeted gene integration and stable expression without disrupting host physiology, represent essential features for model prokaryotic and eukaryotic microbial chassis^49,50^. Despite their importance, neutral loci have not yet been identified or evaluated for gene integration efficiency and expression levels across the *B. methanolicus* genome.

The *in silico* platform CRISPR-COPIES^52^ was employed to identify candidate neutral loci in *B. methanolicus* and design gRNAs for CRISPR-based integration. Utilizing publicly available RNA-sequencing data of *B. methanolicus* under diverse conditions and predicted promoter regions^51^, we selected 12 promising neutral loci characterized by relatively high expression levels of surrounding genes and minimal interference with native promoters for further evaluation (Fig. 4a). A *sfgfp*-expression cassette, driven by the strong constitutive promoter P*_hps_* (the promoter of the *hps* gene encoding Hps, GeneID: BMMGA3_06845), was integrated into these neutral loci. The edited cells exhibited visible green fluorescence due to the high-level expression of sfGFP, confirming successful integration (Fig. 4b). Among the 12 tested loci, 9 demonstrated 100% integration efficiency. Integration efficiencies at NS2, NS4, and NS5 were 58.3%, 10%, and 66.7%, respectively (Fig. 4c). The low integration efficiency at NS4 remains unexplained, as no CDSs or ncRNAs were annotated at this locus.

**Figure 4.**
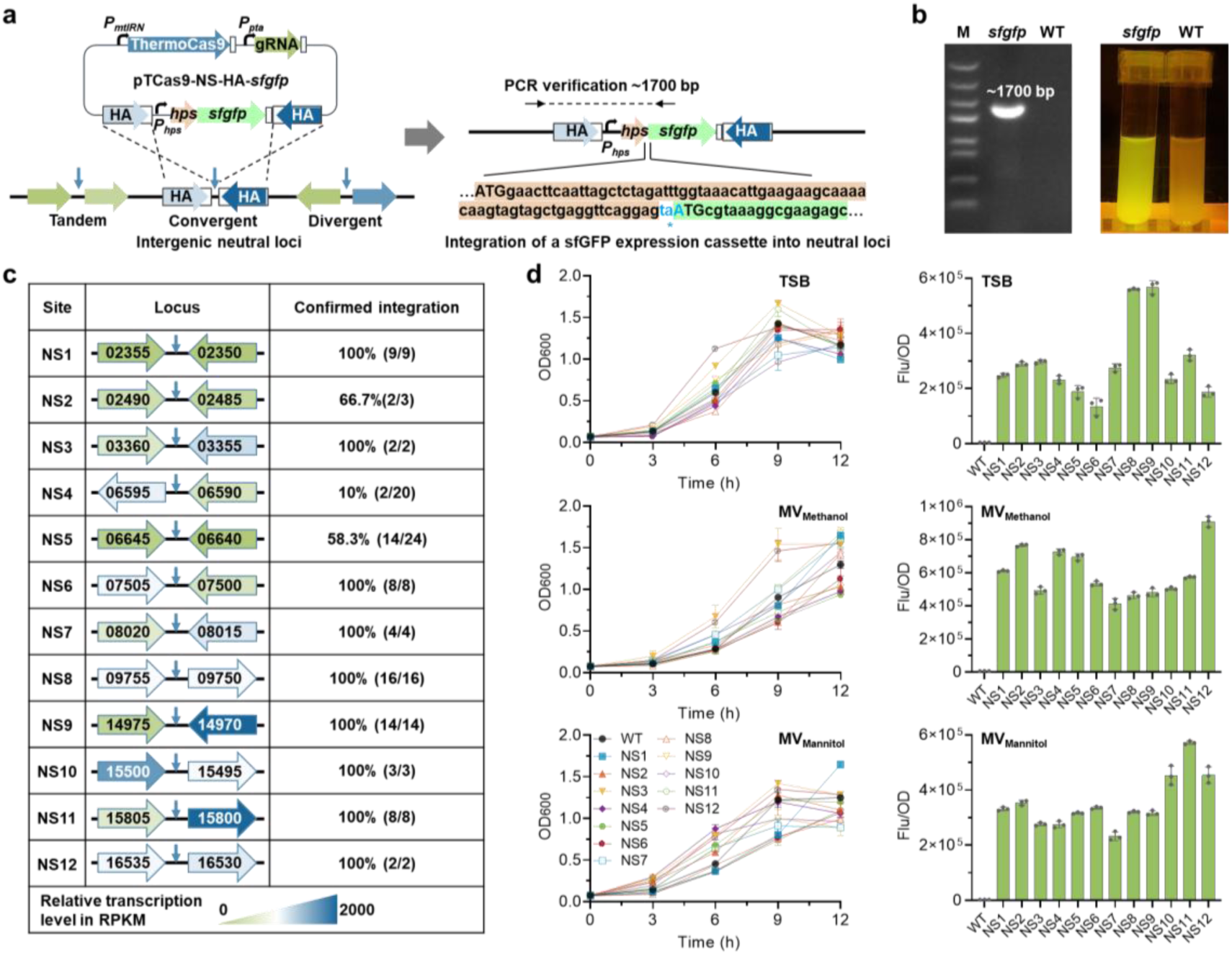
Identification of neutral loci for gene integration and expression. **a,** Schematic representation of the integration of a *sfgfp*-expression cassette into intergenic regions to identify neutral loci. The orientation of surrounding CDSs (convergent, divergent, or tandem) is indicated. **b,** Verification of *sfgfp* integration and expression through PCR and fluorescence visualization. **c,** Integration efficiency of the *sfgfp*-expression cassette at neutral loci, as confirmed by colony PCR of transformants. The orientation of surrounding CDSs is depicted, and the relative transcription levels of these CDSs are visualized using publicly available RNA-sequencing data of *B. methanolicus* under diverse conditions^51^. **d,** Cell growth and sfGFP expression level in strains with the *sfgfp*-expression cassette integrated at neutral loci. MV_Methanol_ and MV_Mannitol_ represent minimal media containing methanol and mannitol as the sole carbon source, respectively. Values and error bars represent the mean ± standard deviation (s.d.) of three biological replicates (n = 3).

We subsequently evaluated cell growth and sfGFP expression in the gene-integrated strains under various conditions. Integration and expression of *sfgfp* across the neutral loci had slight to moderate effects on cell growth, with the exception of NS3 and NS12, where integration enhanced cell growth in all three tested media (TSB rich medium and minimal media with methanol or mannitol as the sole carbon source, respectively) (Fig. 4d). All strains exhibited substantial sfGFP expression across the tested conditions. In TSB rich medium, sfGFP expression levels varied approximately 4-fold among the neutral loci, with the highest expression observed at NS8 and NS9 loci, which showed 422% higher expression compared to the lowest expression at the NS6 locus (Fig. 4d). In minimal media with methanol or mannitol as the sole carbon source, the differences in sfGFP expression levels were less pronounced, with the highest expression at NS12 or NS11 loci being approximately 200% higher than the lowest expression at the NS7 locus (Fig. 4d). Collectively, our study identified several neutral loci suitable for gene integration and expression. The variability in expression levels across different conditions provides a broad range of options for fine-tuning gene expression. A set of ThermoCas9 plasmids containing pre-assembled HAs and validated gRNAs were readily provided, enabling efficient gene integration and expression for diverse experimental and industrial purposes.

### Chromosomal integration of episomal plasmid pBM19 for stable and plasmid-free methylotrophy

*B. methanolicus* is a plasmid-dependent methylotroph, and the curing of the episomal plasmid pBM19 results in the loss of its ability to grow on methanol. Iterative cultivation in methanol-free media has been shown to lead to the loss of methanol-dependent growth in up to 80% of cells^17^. In industrial biomanufacturing, cost-effective complex substrates such as corn steep powder and soybean meal hydrolysate are commonly employed, enabling methanol-independent growth. However, the potential loss of pBM19 during cultivation poses a risk of strain degeneration, which could compromise the efficiency of *B. methanolicus*-based biomanufacturing. Chromosomal integration of the episomal plasmid represents a promising strategy to ensure continuous and stable biosynthesis^53^.

We designed a two-step CRISPR editing strategy to integrate the episomal plasmid pBM19 into the chromosome of *B. methanolicus* ΔpBM69, thereby constructing a plasmid-free methylotrophic strain (Fig. 5a). In the first step, a *sfgfp*-expression cassette flanked by 1 kb HAs, corresponding to the upstream and downstream regions of *repB* (GeneID: BMMGA3_17180, encoding a putative replication initiator protein) on pBM19, was inserted into the neutral locus NS9. In the second step, a ThermoCas9 plasmid expressing two gRNAs targeting *sfgfp* and *repB* was introduced into the *sfgfp*-expressing strain to induce cleavage of both the chromosome and pBM19. Homologous recombination between the chromosome and pBM19 led to the chromosomal integration of pBM19 genes (with *repB* deleted) and the curing of the episomal pBM19 plasmid. Given the potential low efficiency of integrating such large DNA fragments (19 kb) into the chromosome, fluorescence-activated cell sorting (FACS) was employed to screen for edited cells, which lost sfGFP fluorescence due to the replacement of the *sfgfp*-expression cassette with pBM19 genes (Fig. 5b).

**Figure 5.**
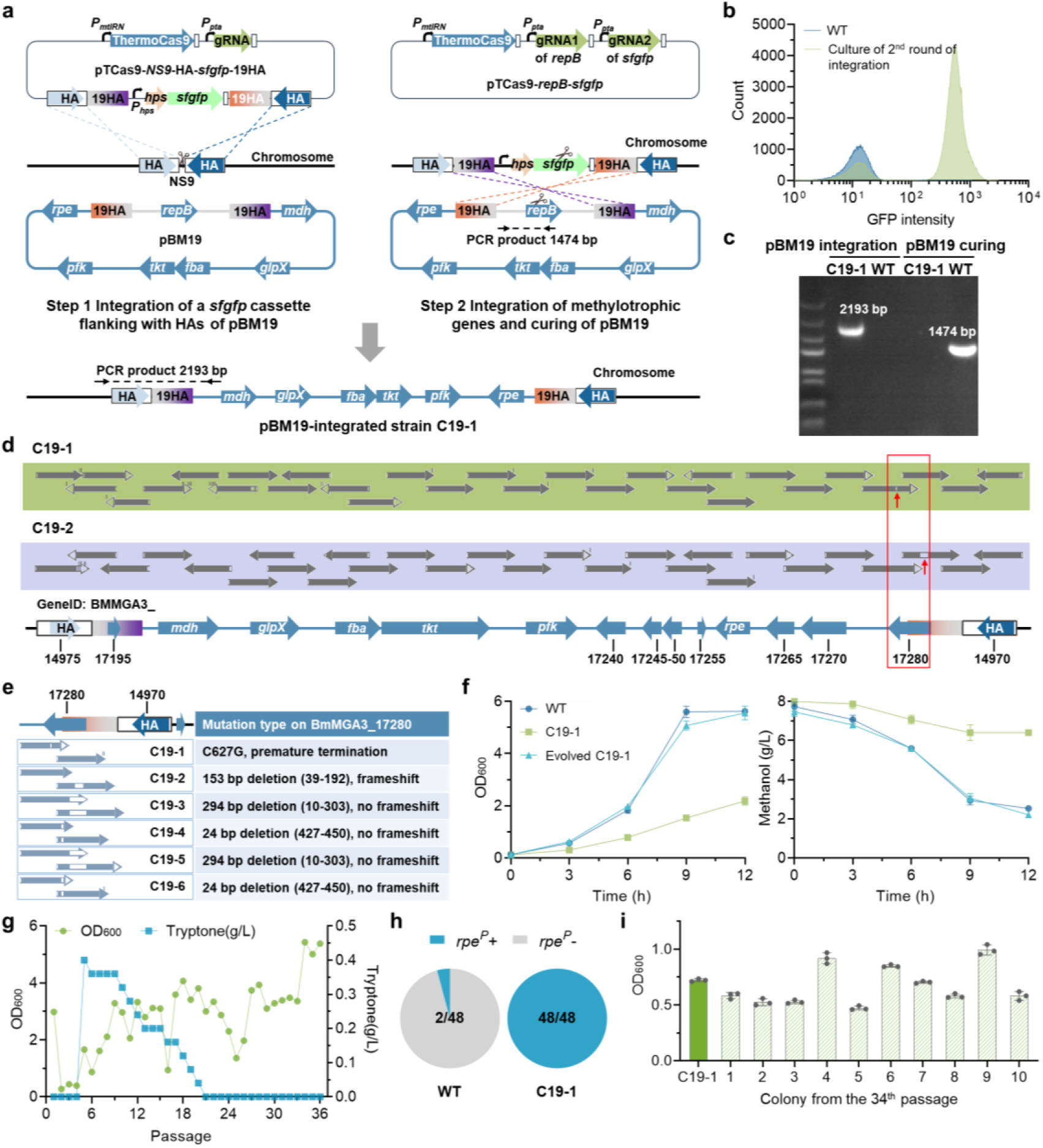
Integration of the large plasmid pBM19 into the chromosome for plasmid-free methylotrophy. **a,** Schematic representation of the two-step CRISPR editing process for chromosomal integration of pBM19. **b,** FACS-based screening of pBM19-integrated cells with the loss of sfGFP fluorescence due to the replacement of the *sfgfp*-expression cassette with pBM19 genes. **c,** Confirmation of chromosomal integration and curing of episomal pBM19 through PCR. **d,** DNA sequencing of the integrated pBM19 genes in two pBM19-integrated strains C19-1 and C19-2. The mutations in the gene BMMGA3_17280 are highlighted in red box and arrows. **e,** Mutations identified in the pBM19 gene BMMGA3_17280 across six pBM19-integrated strains (C19-1 to C19-6). **f,** Cell growth and methanol utilization of the wild-type strain, the original pBM19-integrated strain C19-1, and the evolved C19-1 strain in minimal medium with methanol as the sole carbon source. **g,** ALE of the C19-1 strain in methanol minimal medium, with a gradual reduction in tryptone supplementation. **h,** Stability of pBM19 in *B. methanolicus* wild-type strain and pBM19-integrated strain C19-1 after ten passages of serial cultivation in methanol-free rich medium. PCR of the pBM19-borne *rpe* gene was conducted to assess pBM19 stability. **i,** Growth assay to confirm the stable methylotrophic phenotype of pBM19-integrated strain C19-1. Ten colonies were isolated from the 34^th^ passage of serial cultivation in methanol-free rich medium and then cultivated in minimal medium with methanol as the sole carbon source. Values and error bars represent the mean ± standard deviation (s.d.) of three biological replicates (n = 3).

PCR and DNA sequencing confirmed the successful integration of pBM19 genes at the target chromosomal locus (Fig. 5c, d). Sequencing of the integration region in six edited strains (C19-1 to C19-6) revealed that all strains harbored nonsense mutations, frameshift mutations, or partial deletions in the pBM19 gene BMMGA3_17280, which encodes a hypothetical protein of unknown function (Fig. 5e). The loss-of-function of BMMGA3_17280 appears to facilitate the stable maintenance of pBM19 in the chromosome, although the underlying mechanism remains unclear. The successful chromosomal integration of pBM19 underscores the robust capability of the developed genome editing tool for manipulating large DNA fragments.

The pBM19-integrated strain C19-1 exhibited 11% and 38% lower cell growth rates in TSB rich medium without methanol and methanol minimal medium compared to the wild-type strain, respectively (Supplementary Fig. 7 and Fig. 5f). This reduction in cell growth may explain why pBM19 has not been naturally integrated into the chromosome of *B. methanolicus* during long-term evolution. The copy number of pBM19 in *B. methanolicus* is estimated to range from 10 to 16 copies per chromosome^42^, suggesting that single-copy chromosomal expression of pBM19 genes may be insufficient to support rapid methanol metabolism. Adaptive laboratory evolution (ALE) is a well-established strategy for enhancing cell growth under specific substrate conditions^54^. To improve the growth of the C19-1 strain in methanol minimal medium, we explored the use of ALE. However, cell growth was unstable during the first four serial passages. To facilitate ALE, 0.4 g/L tryptone was added as a nutrient supplement, and its concentration was gradually reduced to zero (Fig. 5g). Ultimately, after 34 passages, the cell growth of strain C19-1 in minimal medium with methanol as the sole carbon source was restored to levels comparable to the wild-type *B. methanolicus* strain (Fig. 5f).

To assess the stability of methylotrophy, the wild-type strain harboring pBM19 and the pBM19-integrated strain C19-1 were iteratively cultivated in methanol-free media. After 10 passages (∼40 generations), 46 out of 48 tested colonies (∼96%) of the wild-type strain lost the pBM19 plasmid. In contrast, all 48 tested colonies of the C19-1 strain retained the pBM19 genes (Fig. 5h and Supplementary Fig. 8). Even after 34 passages (∼140 generations), all tested colonies of the C19-1 strain maintained the pBM19 genes and the ability to grow in methanol minimal medium (Fig. 5i). These results demonstrate that single-copy chromosomal expression of pBM19 genes can support efficient methylotrophy. The engineered strain C19-1 represents a plasmid-free methylotrophic model with stable and robust growth, free from the risk of plasmid loss and strain degeneration.

### A cloud-based GEM enabling native and heterologous pathway design and visualization

GEMs that mathematically describe gene-protein-reaction relationships serve as fundamental tools for the DBTL cycle in synthetic biology^23^. Despite the widespread applications, no GEM has been developed for the thermophilic methylotroph *B. methanolicus*. To address this gap, we constructed the first GEM *i*BM822 through an iterative optimization pipeline (Fig. 6a). The draft model was initially built using CarveMe with the BiGG database as a foundational framework^55^. The draft model was subsequently refined through iterative manual curation that integrated KEGG and MetaCyc databases reactions, followed by automated method to remove errors in the model^56,57^. Critical improvements included experimental determination of amino acid composition for biomass equation refinement (Supplementary Table 1). The final model, designated *i*BM822, comprises 822 protein-coding genes, 1,479 metabolites, and 1,730 compartmentalized reactions (cytosolic, periplasmic, extracellular). The MEMOTE^58^ comprehensive score of *i*BM822 is 93 (Supplementary Fig. 9), surpassing established models like *E. coli i*ML1515 (score 91)^59^, *B. subtilis i*YO844 (score 86)^60^, and *C. glutamicum i*CW773 (score 21)^61^.

**Figure 6.**
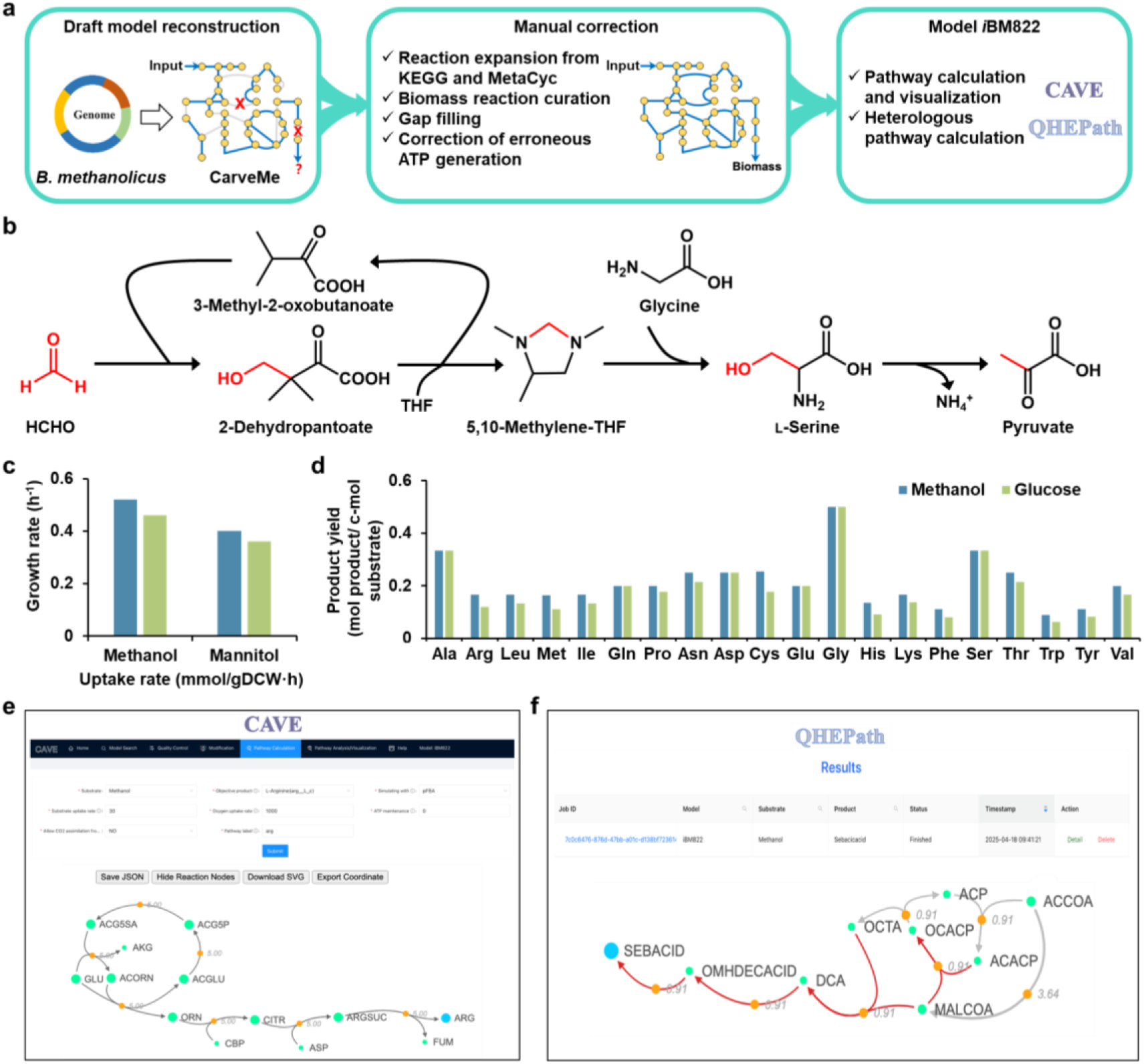
Construction and application of *B. methanolicus* GEM *i*BM822. **a,** Iterative construction pipeline of GEM *i*BM822. **b,** A predicted formaldehyde assimilation pathway via 5,10-methylene-THF. **c,** Comparison of calculated growth rates with experimental values for different carbon sources. Methanol and mannitol were utilized at rate of 30 mmol/gDCW·h and 4.8 mmol/gDCW·h, respectively. Data of substrate uptake rate and specific cell growth rate reported by Delépine were used for comparison^62^. **d,** Theoretical amino acid yields from methanol versus glucose substrates calculated using *i*BM822. **e,** Visualization of methanol-to-L-arginine biosynthesis using *i*BM822-integrated CAVE^63^. **f,** *De novo* design of sebacic acid pathway using *i*BM822-integrated QHEPath^64^. The four heterologous reactions are highlighted in red arrows.

Beyond the well-characterized ribulose monophosphate (RuMP) cycle, *i*BM822 revealed a putative formaldehyde assimilation route via 5,10-methylenetetrahydrofolate (5,10-methylene-THF) (Fig. 6b). Thermodynamic analysis showed two spontaneous reactions (ΔG = −19.2 and −6.3 kJ/mol, respectively) catalyzed by ketopantoate hydroxymethyltransferase (BMMGA3_10620), which have been detected in other organisms^65,66^:

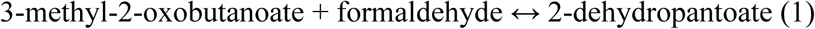

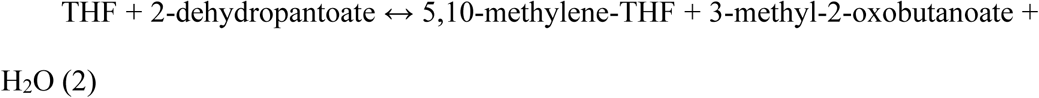

The resulting 5,10-methylene-THF could theoretically combine with glycine through glycine hydroxymethyltransferase (BMMGA3_16040) to yield L-serine, which is subsequently deaminated to pyruvate by L-serine deaminase (BMMGA3_05935 and BMMGA3_05940). However, gene knockout experiments demonstrated that disrupting key RuMP pathway genes *hps* or *phi* completely abolished methanol-dependent growth, suggesting this alternative pathway may not be physiologically active under standard conditions.

To assess predictive accuracy of *i*BM822, we simulated growth on methanol and mannitol – the two primary carbon sources for *B. methanolicus*. The computed growth rates (0.52 h^−1^ and 0.4 h^−1^, respectively) exceeded experimental values^62^ by 11–13% (Fig. 6c), indicating robust model performance. Given the industrial relevance of methanol-based amino acid production^67^, we compared theoretical yields of 20 proteinogenic amino acids from methanol versus glucose. Fourteen amino acids showed 13–50% higher methanol-derived yields, while six (L-alanine, L-glutamine, L-aspartate, L-glutamate, glycine, and L-serine) exhibited equivalent yields from both substrates (Fig. 6d). This analysis supports the preference of methanol as a feedstock for sustainable amino acid production.

For the convenience of researchers without coding expertise, GEM *i*BM822 was integrated into the two cloud-based platforms. CAVE (https://cave.biodesign.ac.cn/)^63^: A cloud-based interface enabling non-expert users to visualize metabolic pathways and fluxes and perform reaction editing including modifying the fluxes of reactions and manually adding heterologous reactions. An example of calculating and visualizing the L-arginine biosynthetic pathway is shown (Fig. 6e). QHEPath (https://qhepath.biodesign.ac.cn/)^64^: A heterologous pathway design module that automatically integrates foreign reactions into *i*BM822. We demonstrated this functionality by designing a sebacic acid (a monomer for nylon) pathway requiring four heterologous steps (Fig. 6f), highlighting its potential for biodesign of non-native chemical synthesis.

### Characterizing the feedback regulation of L-arginine biosynthesis for L-arginine overproduction from methanol

L-Arginine, the only proteinogenic amino acid featuring a guanidino side chain, has extensive applications in the pharmaceutical, nutraceutical, and feed industries. It plays critical roles in stimulating growth hormone secretion, enhancing endogenous nitric oxide metabolism, and promoting wound healing^68^. Currently, L-arginine is predominantly produced using engineered strains of *E. coli* or *Corynebacterium glutamicum*, with glucose as the primary carbon source^69,70^. To address the growing market demand and achieve sustainable amino acid production, methanol has emerged as a promising alternative feedstock^71,72^. However, to date, no studies have reported the production of L-arginine from methanol.

Amino acid biosynthesis is typically tightly regulated through feedback inhibition at transcriptional and/or enzymatic levels^72^. The L-arginine biosynthesis pathways and feedback inhibition mechanisms in *E. coli* and *C. glutamicum* have been extensively characterized^69,70^. In *E. coli*, L-arginine biosynthesis proceeds via a linear pathway initiated by acetylglutamate synthase (ArgA), which is subject to feedback inhibition by high intracellular L-arginine concentrations^69^. In contrast, *C. glutamicum* employs an acetyl-CoA recycling pathway, in which a bifunctional L-glutamate and L-ornithine acetyltransferase (ArgJ) catalyzes the formation of acetylglutamate (Fig. 7a). Acetylglutamate kinase (ArgB), the key rate-limiting enzyme in this pathway, is feedback-inhibited by L-arginine^70^. GEM simulation suggests that *B. methanolicus* shares a similar acetyl-CoA recycling pathway with *C. glutamicum* (Fig. 6e and Fig. 7a), whereas its regulatory mechanisms remain unexplored. To engineer *B. methanolicus* for methanol-based L-arginine production, it is essential to characterize and alleviate these regulatory constraints.

**Figure 7.**
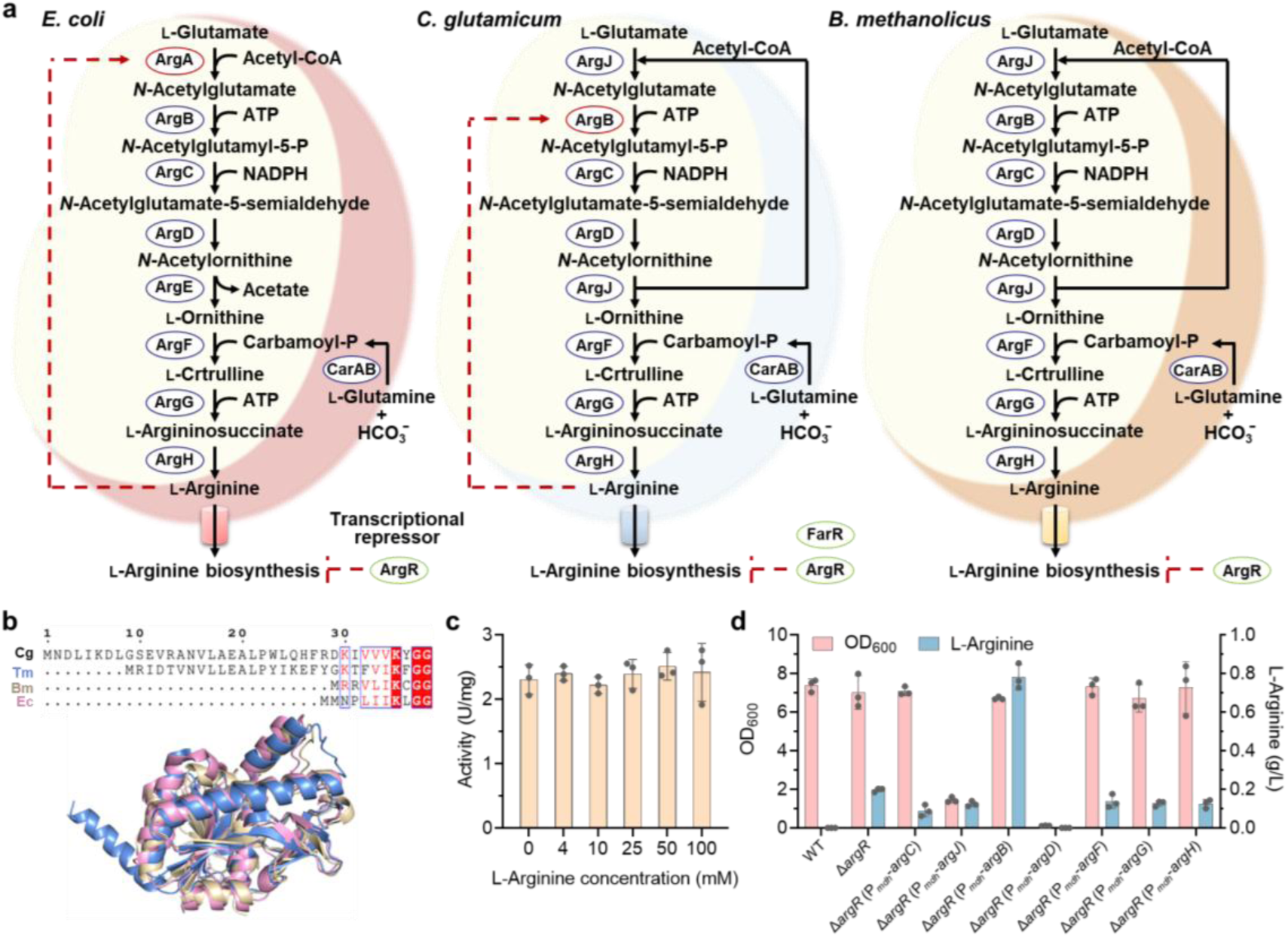
L-Arginine biosynthesis is regulated at the transcriptional level but not at the enzyme level in *B. methanolicus*. **a,** Schematic representation of different L-arginine biosynthetic pathways and regulatory mechanisms in microorganisms. Red arrows and lines indicate feedback inhibition of enzymes and transcriptional repression by L-arginine, respectively. Enzymes highlighted in red ovals are subject to feedback inhibition by L-arginine. ArgA: acetylglutamate synthase; ArgB: acetylglutamate kinase; ArgC: acetyl-glutamyl-phosphate reductase; ArgD: acetylornithine aminotransferase; ArgE: acetylornithine deacetylase; ArgF: ornithine carbamoyltransferase; ArgG: argininosuccinate synthase; ArgH: argininosuccinate lyase; ArgJ: bifunctional glutamate and ornithine acetyltransferase; ArgR and FarR, L-arginine repressor; CarAB, carbamoyl phosphate synthetase. **b,** Sequence and structure alignment of ArgB proteins from *B. methanolicus* (BmArgB, golden), *E. coli* (EcArgB, magenta), and *Thermotoga maritima* (TmArgB, blue). The structure of BmArgB was predicted using AlphaFold2^73^ and aligned with the experimentally determined structures of EcArgB (PDB: 1GS5)^74^ and TmArgB (PDB: 2BTY)^75^. **c,** Effects of L-arginine addition on the catalytic activity of ArgB from *B. methanolicus*. **d,** Effects of deletion of *argR* and overexpression of L-arginine biosynthetic genes on the accumulation of extracellular L-arginine. Values and error bars represent the mean ± standard deviation (s.d.) of three biological replicates (n = 3).

The crystal structure of L-arginine-complexed ArgB from *Thermotoga maritima* reveals a ring-like hexamer formed by the association of three homodimer subunits through an interlaced N-terminal α-helix. Binding of L-arginine at the inter-dimeric junction stabilizes an enlarged active site conformation, which impedes catalysis and mediates feedback inhibition of ArgB by L-arginine^75^. In contrast, the L-arginine-insensitive ArgB from *E. coli* adopts a homodimeric structure and lacks both the N-terminal α-helix and the L-arginine binding site^74^. Sequence and structural alignments indicate that *B. methanolicus* ArgB (GeneID: BMMGA3_04430) also lacks the N-terminal α-helix, resembling the L-arginine-insensitive ArgB of *E. coli* and differing from the L-arginine-sensitive ArgBs of *C. glutamicum* and *T. maritima* (Fig. 7b). To confirm the L-arginine insensitivity of *B. methanolicus* ArgB, the enzyme was heterologously expressed and purified (Supplementary Fig. 10). No inhibition of catalytic activity was observed in the presence of 100 mM L-arginine (Fig. 7c), consistent with the behavior of the L-arginine-insensitive *E. coli* ArgB^74^.

Although *B. methanolicus* ArgB is not subject to feedback inhibition, the wild-type strain is incapable of overproducing L-arginine, indicating that L-arginine biosynthesis is regulated at other levels. In *E. coli*, the transcriptional repressor ArgR negatively regulates the expression of L-arginine biosynthetic genes^69^. In *C. glutamicum*, two transcriptional repressors, ArgR and FarR, act synergistically to control L-arginine biosynthesis^70^. BLAST analysis identified a putative ArgR in *B. methanolicus* (GeneID: BMMGA3_11380), which shares 27% and 30% amino acid sequence similarity with the ArgR proteins of *E. coli* and *C. glutamicum*, respectively. No homolog of *C. glutamicum* FarR was detected in the *B. methanolicus* genome. To investigate its role, *argR* was deleted in *B. methanolicus* using the CRISPR editing tool.

The *B. methanolicus* Δ*argR* mutant produced and excreted approximately 0.2 g/L L-arginine during shake flask fermentation with methanol as the sole carbon source (Fig. 7d), demonstrating that the repression of L-arginine biosynthesis was alleviated by *argR* deletion. In summary, L-arginine biosynthesis in *B. methanolicus* is regulated solely at the transcriptional level, in contrast to the dual-level (transcriptional and enzymatic) regulation observed in *E. coli* and *C. glutamicum*^59,60^. To identify the bottleneck of the L-arginine biosynthetic pathway, genes of the *argCJBDF* and *argGH* operons were individually overexpressed in Δ*argR* mutant. *argB* overexpression largely enhanced L-arginine titer to approximately 0.8 g/L, a 4-fold improvement compared to Δ*argR*. Overexpression of other genes brought negative effects on L-arginine production or dramatically impaired cell growth (Fig. 7d). The results suggest that although ArgB is not feedback inhibited by L-arginine, it is the rate-limiting step for L-arginine biosynthesis.

## Discussion

Methylotrophs play pivotal roles in global biogeochemical processes^4–6^ and have recently gained significant attention for their applications in the biomanufacturing of chemicals, fuels, and foods from C1 feedstocks^7–10^. However, compared to model and industrial chassis such as *E. coli* and *S. cerevisiae*, synthetic biology tools for methylotrophs have historically been limited in both sophistication and scope, hindering both fundamental research and engineering applications. Recent advancements in synthetic biology technologies have significantly expanded the potential for engineering methylotrophs. A notable example is the methylotrophic yeast *Komagataella* (*Pichia*) *pastoris*, which has been extensively utilized for industrial enzyme and vaccine production. The integration of advanced CRISPR tools and metabolic modeling has further broadened its product portfolio to include various secondary metabolites, such as plant natural products, fatty acids, and their derivatives^76–79^.

In contrast to methylotrophic yeasts like *K. pastoris*, *B. methanolicus* is a methylotrophic bacterium and an extremophile, characterized by rapid growth at high temperatures (50–55°C; doubling time ∼1.1 h) and adaptability to seawater-based media^20^. Next-generation industrial biotechnology (NGIB) leveraging extremophiles such as *B. methanolicus* enables cost-effective biomanufacturing with reduced energy requirements for sterilization and cooling^19^. With its robust primary metabolism, *B. methanolicus* holds significant potential for the production of amino acids, vitamins, alcohols, organic acids, and feed proteins^19,70–73^.

To fully harness the potential of *B. methanolicus* for diverse applications, we developed a ready-to-use synthetic biology toolkit for the research community. This toolkit includes a simple and efficient electroporation protocol, a robust CRISPR-based method for multiplex genome editing, neutral loci for tunable overexpression of integrated genes, and a GEM compatible with cloud-based visualization and heterologous pathway design. Similar toolkits are plentifully available in *E. coli*, where they have significantly strengthened capacities for genetic engineering^80–82^. We anticipate that our toolkit will similarly expand the engineering possibilities for *B. methanolicus*, complementing other recently developed constitutive and inducible promoters^26,39,40^ and compatible plasmids^39^.

Additionally, the application of our toolkit enabled the creation of a novel plasmid-free chassis, C19-1, through the curing of pBM69 and chromosomal integration of pBM19. Given its stable and robust methylotrophic growth without degeneration, C19-1 holds great promise for elucidating the evolutionary mechanisms of plasmid-dependent methylotrophy and developing methylotrophic cell factories for biomanufacturing. Previous studies have demonstrated that *E. coli-B. methanolicus* shuttle plasmids pNW33N (with the pC194 replicon from *Staphylococcus aureus*) and pHP13 (with the pTA1060 replicon from *B. subtilis*) are compatible in wild-type *B. methanolicus* harboring pBM19 and pBM69^39^. Consequently, the curing of pBM19 and pBM69 frees up two additional replicons (pBM19 and pBM69) for the development of compatible plasmids, enabling the stable maintenance of up to four plasmids in strain C19-1.

We employed our toolkit to address a key question regarding this plasmid-dependent methylotroph: the physiological function of chromosomal copies of methylotrophic genes (*mdh2*, *mdh3*, *act*, *rpe*, *hps*, *phi*, *pfk*, *glpX*, *fba*, and *tkt*) in methanol metabolism. With the exception of *act*, *hps*, and *phi*, which exist exclusively on the chromosome, the remaining genes have an additional copy on the pBM19 plasmid (*mdh*^P^, *rpe*^P^, *pfk*^P^, *glpX*^P^, *fba*^P^, and *tkt*^P^). Transcription of pBM19 genes is typically induced by methanol, whereas chromosomal copies are not, leading to the hypothesis that the chromosomal copies are dispensable for methylotrophy^15^. Although enzymes encoded by chromosomal and plasmid copies often exhibit distinct catalytic and biochemical properties^45,46,83–86^, there remains a gap in understanding their physiological roles. CRISPR-enabled genome editing and growth assays of gene-deletion mutants have provided critical insights to bridge this gap. By integrating previous *in vitro* findings with our *in vivo* results, we conclude that several chromosomal gene copies, including *rpe*, *pfk*, *hps*, and *phi* are essential for methylotrophic growth. *pfk* is also indispensable for mannitol metabolism.

A hallmark of synthetic biology is the integration of computer-assisted design with genome editing to engineer cells with specific functionalities, for which GEMs serve as indispensable tools^87^. The first GEM for *B. methanolicus i*BM822 is presented here. However, the utilization and customization of such computational tools typically require researchers to possess a certain level of coding proficiency. To make these tools accessible, particularly for researchers without coding expertise, the GEM *i*BM822 was integrated into two user-friendly platforms: CAVE^63^ and QHEPath^64^. CAVE enables users to easily add, remove, or modify biochemical reactions within the model for pathway design and visualization. QHEPath facilitates the automatic incorporation of heterologous reactions into the model, enabling the design of pathways with maximal yield. With the recent advancements in large language models (LLMs), LLM-empowered metabolic modeling is anticipated to provide novel metabolic insights and precise predictions for bioengineering applications^88^. A preliminary analysis using these biodesign tools highlighted the advantages of *B. methanolicus* for amino acid production from methanol compared to traditional glucose-based biosynthesis. As a first step toward engineering *B. methanolicus* for methanol-based amino acid biomanufacturing, we applied these tools to characterize and release feedback regulation of L-arginine biosynthesis. Although the current L-arginine production level represents an early-stage prototype, further synthetic biology-guided metabolic engineering is expected to yield efficient methylotrophic cell factories suitable for industrial-scale applications.

## Methods

### Strains and culture conditions

The strains utilized in this study are enumerated in Supplementary Table 2. For plasmid cloning, *E. coli* strains Trans1-T1 (*dam*^+^) and Trans110 (*dam*^−^) (TransGen Biotech, Beijing, China) were aerobically cultivated at 37°C in Luria-Bertani (LB) broth. The medium was supplemented with chloramphenicol (20 μg/mL) or kanamycin (50 μg/mL) as necessary. *B. methanolicus* and its derivatives were cultured in 500 mL baffled shake flasks containing 50 mL of TSB medium^89^ or minimal medium^32^, with shaking at 220 rpm and at 50°C. Methanol (250 mM) or mannitol (30 mM) was added to the minimal medium as the carbon source, with chloramphenicol (5 μg/mL) added when required. For evaluation of neutral loci for sfGFP expression, the engineered strains were inoculated into TSB medium for pre-cultivation. Subsequently, the culture was inoculated at an initial OD_600_ of 0.1 into a 96-well deep-well plate containing 800 μL of TSB or minimal medium (with 250 mM methanol or 30 mM mannitol). Cultivation was conducted at 50°C with shaking at 800 rpm. Samples were collected periodically and determined for OD_600_ and sfGFP fluorescence. For L-arginine production in shake flasks, *B. methanolicus* and its derivatives were cultivated in 500 mL shake flasks containing 50 mL of TSB medium. The culture served as a seed to inoculate 50 mL of minimal medium containing methanol (250 mM) as the carbon source in 500 mL shake flasks. Strains were cultivated with shaking at 220 rpm and at 50°C. Samples were collected periodically and determined for OD_600_ and L-arginine titer.

### Plasmid construction

The plasmids and primers employed in this study are detailed in Supplementary Table 2 and Supplementary Table 3, respectively. Plasmid construction was achieved through recombination using the ClonExpress MultiS One Step Cloning Kit (Vazyme, Nanjing, China). DNA polymerase and PCR reagents were procured from TransGen Biotech (Beijing, China), while restriction endonucleases were obtained from New England Biolabs (Beijing) (Beijing, China). DNA synthesis and sequencing services were rendered by GENEWIZ Inc. (Suzhou, China). The comprehensive procedure for plasmid construction is delineated in Supplementary Table 4. The nucleotide sequence of the pTCas9-Goi-HA plasmid (with *amyX* deletion as an example) is presented in Supplementary Table 5.

### Electroporation of *B. methanolicus*

The optimization of competent cell preparation and electroporation was based on the protocol developed by Jakobsen et al.^17^. The initial optimization step involved the elimination of PEG8000 from the EP washing buffer, resulting in a new washing buffer containing solely 1 mM HEPES at pH 7.0. Subsequent steps included testing PEGs of varying molecular weights (PEG2000, PEG4000, and PEG6000) to replace PEG8000 in the EP resuspension buffer (1 mM HEPES, 25% PEG8000, pH 7.0), altering the PEG8000 concentration in the EP resuspension buffer (15%, 20%, 25%, and 35%), optimizing the recovery time post-electroporation (from the original two-passage recovery of 16 h and 6 h to a single-passage recovery of 3 h, 6 h, and 12 h), adjusting the OD_600_ at which cells were harvested for competent cell preparation (from 0.25 to 0.5 and 0.7), and testing varying amounts of plasmid DNA for electroporation (0.2 μg, 0.5 μg, 1 μg, and 2 μg). Post-electroporation and recovery, cells were plated on TSB solid medium supplemented with chloramphenicol and incubated at 50°C for 24–36 h until colony formation. For routine plasmid transformation, plasmids were prepared in *E. coli* Trans1-T1 (*dam*^+^), resulting in methylated plasmids. To assess the impact of DNA methylation on transformation efficiency, methylation-free plasmids were prepared in *E. coli* Trans110 (*dam*^−^).

### Evaluation of promoter strength

The fluorescent reporter sfGFP was employed to assess promoter strength^38^. A *sfgfp*-expression cassette, regulated by the target promoter, was assembled into the pNW33N plasmid (GenBank: AY237122.1). The recombinant plasmid was introduced into *E. coli* and *B. methanolicus*, with transformants verified via colony PCR and subsequently cultured for fluorescence determination. For xylose- and mannitol-inducible sfGFP expression, xylose (3.3 mM) and mannitol (2.5 mM) were added as inducers, as previously described^39^. Fluorescence outputs were detected using cells in the stationary growth phase.

### CRISPR-mediated genome editing in *B. methanolicus*

Routine CRISPR-mediated gene deletion in *B. methanolicus* involved the transformation of 1 μg of the ThermoCas9 plasmid, harboring ThermoCas9, a targeting gRNA, and two HAs, via electroporation. For gene integration, the gene of interest, flanked by two HAs, was assembled into the ThermoCas9 plasmid. For the curing of endogenous plasmids pBM19 and pBM69, HAs were unnecessary, and only ThermoCas9 and a plasmid-targeting gRNA were expressed in the ThermoCas9 plasmid. Post-electroporation and recovery, cells were plated on TSB solid medium supplemented with chloramphenicol and incubated at 50°C for 24–36 h. Colonies were verified by PCR, and those with the correct PCR profile were selected and cured of the ThermoCas9 plasmid.

For plasmid curing, colonies were cultured in TSB medium without chloramphenicol but with mannitol at 58°C overnight, then transferred into fresh medium for a second passage. Mannitol was added to enhance ThermoCas9 expression, intended to impose metabolic burden and inhibit cell growth, thereby favoring cells that had lost the ThermoCas9 plasmid. Cells were then plated on TSB solid medium without chloramphenicol and mannitol. Colonies were confirmed as cured by PCR amplification of a specific fragment of the ThermoCas9 plasmid and by determining their chloramphenicol sensitivity. Transformants with cured plasmids were utilized for subsequent genome editing or other tests. All plasmids and gRNAs used for *B. methanolicus* genome editing are listed in Supplementary Table 2 and Supplementary Table 6, respectively.

### Chromosomal integration of endogenous large plasmids

A two-step CRISPR editing process was employed for the chromosomal integration of pBM19 into *B. methanolicus* ΔpBM69. The first step involved integrating a *sfgfp*-expression cassette, flanked by 1 kb HAs homologous to the upstream and downstream regions of *repB* in pBM19, into the neutral locus NS9, following the standard CRISPR-mediated genome editing procedure in *B. methanolicus*. After curing the first ThermoCas9 editing plasmid, the engineered strain was used for competent cell preparation. Subsequently, a second ThermoCas9 editing plasmid, expressing ThermoCas9 and two *sfgfp*- and *repB*-targeting gRNAs, was transformed into the competent cells. The second step entailed the cleavage of both the chromosome and pBM19, followed by homologous recombination between the pre-set HAs at the neutral locus NS9 and pBM19. Post-chromosomal integration of pBM19, *repB* in pBM19 and *sfgfp* at the neutral locus NS9 were excised. Consequently, FACS was employed to select pBM19-integrated cells that had lost sfGFP fluorescence. Cells were harvested by centrifugation at 6,000 × *g* for 10 min, washed once, and resuspended in phosphate buffer (pH 7.4). sfGFP fluorescence was analyzed by flow cytometry (MoFlo XDP, Beckman Coulter, USA) with the following parameters: excitation at 488 nm, emission fluorescence at 529 ± 14 nm, sample pressure of 60 psi, and a nozzle diameter of 70 μm. Cells were captured on the signal channels of FITC (voltage 510 V), FSC (voltage 150 V), and SSC (voltage 250 V). All captured events were used for fluorescence analysis, with data analyzed using Beckman Summit software v5.2. Cells devoid of sfGFP fluorescence were collected and plated on TSB solid medium. Colony PCR was used to verify the curing of the pBM19 plasmid and the chromosomal integration of pBM19 genes.

### Complementation of pBM19 in *B. methanolicus* using pBMEC22

To complement pBM19 in the *B. methanolicus* ΔpBM19 strain, pBM19 was extracted from the *B. methanolicus* wild-type strain using the Gram-Positive Bacterium Plasmid Extraction Mini Kit (Solarbio, Beijing, China). pBM19 was linearized using *Sac*I. The pSC101 origin and kanamycin resistance gene were amplified by PCR from the pSB4K5 plasmid (GenBank: EU496099.1) using the primer pair pSB-F/pSB-R. The PCR products were purified and ligated with linearized pBM19 via recombination using the ClonExpress MultiS One Step Cloning Kit (Vazyme, Nanjing, China). The resultant 22 kb *E. coli*-*B. methanolicus* shuttle plasmid, designated pBMEC22, was transformed into the *B. methanolicus* ΔpBM19 strain via electroporation. As only transformants harboring pBMEC22 could proliferate in methanol minimal medium, the recovery culture was transferred to methanol minimal medium for the selection of correct transformants. Cells were then plated on TSB solid medium to isolate single colonies, which were verified by PCR and growth tests in methanol minimal medium.

### Construction and refining of the GEM

The GEM *i*BM822 was constructed using the genomic data of *B. methanolicus* MGA3 (RefSeq: GCF_000724485.1) via the CarveMe pipeline, an automated Python-based framework for metabolic model reconstruction^55^. Temporary gene IDs (e.g. WP_003348775.1) from NCBI were mapped to *B. methanolicus* specific identifiers (e.g. BMMGA3_05340) using BLASTp against Uniprot genome annotation sequences (e-value ≤ 10−5, identity ≥ 80%). To improve completeness, the draft model was manually curated by supplementing missing genes, reactions, and metabolites based on information from the MetaCyc and KEGG databases. Biomass composition (DNA, RNA, lipids, peptidoglycan, carbohydrates, and small molecules) was defined based on published literature of *B. subtilis* 168 model *i*YO844^60^. The composition of macromolecular protein in *B. methanolicus* was experimentally determined, with the detailed amino acid composition provided in Supplementary Table 1. Metabolic gaps were filled using a weighted parsimonious FBA (pFBA) algorithm for identification of non-synthesized biomass composition with the cross-species metabolic network model CSMN built by BiGG^64^. To address erroneous ATP generation, reaction reversibility was automatically corrected using the workflow established during the construction of the CSMN model^64^. The modified reactions were subsequently manually validated by cross-referencing MetaCyc, KEGG, and thermodynamic data from eQuilibrator. *In silico* analyses of cell growth and amino acid biosynthesis using different carbon sources were performed using the CAVE platform^63^ integrated with the GEM *i*BM822. Prediction of sebacic acid biosynthetic pathway was performed using the QHEPath platform^64^ integrated with the GEM *i*BM822.

### Heterologous expression, purification, and enzyme activity assay of ArgB

The *argB* gene was amplified from the genomic DNA of *B. methanolicus* using the primer pair argB-21a-F and argB-21a-R. The PCR product was purified and ligated with pET-21a, which had been digested with *Nde*I and *Xho*I. ArgB fusing to a C-His tag was expressed in *E. coli* BL21 (DE3) and purified using His Spin Trap columns (GE Healthcare, USA). Protein concentration was determined using the Pierce^TM^ BCA Protein Assay kit (Thermo Scientific, USA). Purified proteins were analyzed on a 15% SurePAGE^TM^ gel (GenScript, China). The enzyme activity of purified ArgB was determined using the method described by Haas et al.^90^, with the enzymatic reaction conducted at 50°C.

### Analytical methods

The sfGFP fluorescence of cells in the stationary growth phase was determined using a microplate reader (Infinite E Plex CN/30223586, Tecan Life Sciences, Switzerland), with excitation and emission wavelengths set at 488 nm and 520 nm, respectively. Fluorescence signals were normalized by OD_600_ values. Methanol in the medium was quantified using the M-100 Biosensor Analyzer (Sieman Technology, Shenzhen, China), equipped with an alcohol oxidase membrane. The analytical signal was derived from the quantification of H_2_O_2_ production, generated by methanol oxidation catalyzed by alcohol oxidase^91^. Mannitol in the medium was determined using High Performance Liquid Chromatography (HPLC) (Agilent 1260 Infinity HPLC, Agilent Technologies, USA), equipped with a RID detector and an Aminex HPX-87H column (300 mm × 7.8 mm, 9 µm, Bio-Rad Laboratories, USA). The analysis was performed with a mobile phase of 5 mM H_2_SO at 55°C and a flow rate of 0.5 mL/min^91^. Amino acid concentrations in the fermentation broth were determined using HPLC, equipped with a UV detector and a ZORBAX Eclipse AAA column (4.6 mm × 150 mm, 5 μm, Agilent Technologies, USA). A gradient of 50 mM sodium acetate buffer at pH 6.4, with a gradient solution containing acetonitrile-water (50%, v/v), was used as the eluent. Amino acids were detected as their 2,4-dinitrofluorobenzene derivatives at 360 nm, following the precolumn derivation method^24^.

### Data availability

The data supporting the findings of this work are available within the paper and the Supplementary Information files. The files for GEM *i*BM822 are available from github at https://github.com/YuWangLab/iBM822.

## Competing interests

The authors declare no competing financial interests.

## Author contributions

Y.W. conceived and designed this project. P.L., X.Y., Q.W., T.C., Y.B., T.Z., J.Y., and S.G. performed the experiments. Q.Y., P.W., and H.M. constructed the metabolic model and performed *in silico* analyses. P.L., C.X. Z.Z., B.X., and Y.W. analyzed the data. C.X., Z.Z., B.X., H.M., and Y.W. contributed reagents and analytic tools. Y.W. supervised the research and wrote the initial paper draft. All authors contributed to discussion and writing of the final paper.

## Acknowledgments

This work was supported by the National Key R&D Program of China (2024YFA0918100), the National Natural Science Foundation of China (32222004, 32400069, 32401272, and 32470057), the Tianjin Science Fund for Distinguished Young Scholars (24JCJQJC00010), the China Postdoctoral Science Foundation (2024M753446 and 2024M753447), the Tianjin Synthetic Biotechnology Innovation Capacity Improvement Project (TSBICIP-KJGG-019 and TSBICIP-PTJJ-011), and the Youth Innovation Promotion Association of Chinese Academy of Sciences (2021177).

## Notes

### Competing Interest Statement

The authors have declared no competing interest.

