## Supplementary Fig. 1-10 for "A synthetic biology toolkit for interrogating plasmid-dependent methylotrophy and enhancing methanol-based biosynthesis of *Bacillus methanolicus*"

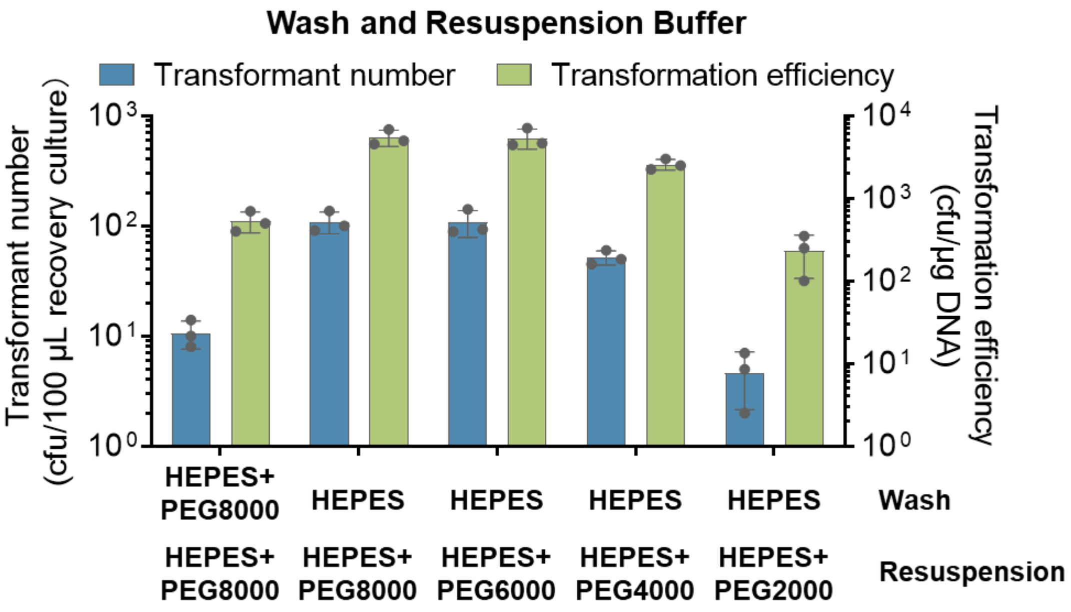


**Supplementary Fig. 1.** Optimization of wash and resuspension buffer for electro-transformation. Values and error bars represent the mean ± standard deviation (s.d.) of three biological replicates (n = 3).

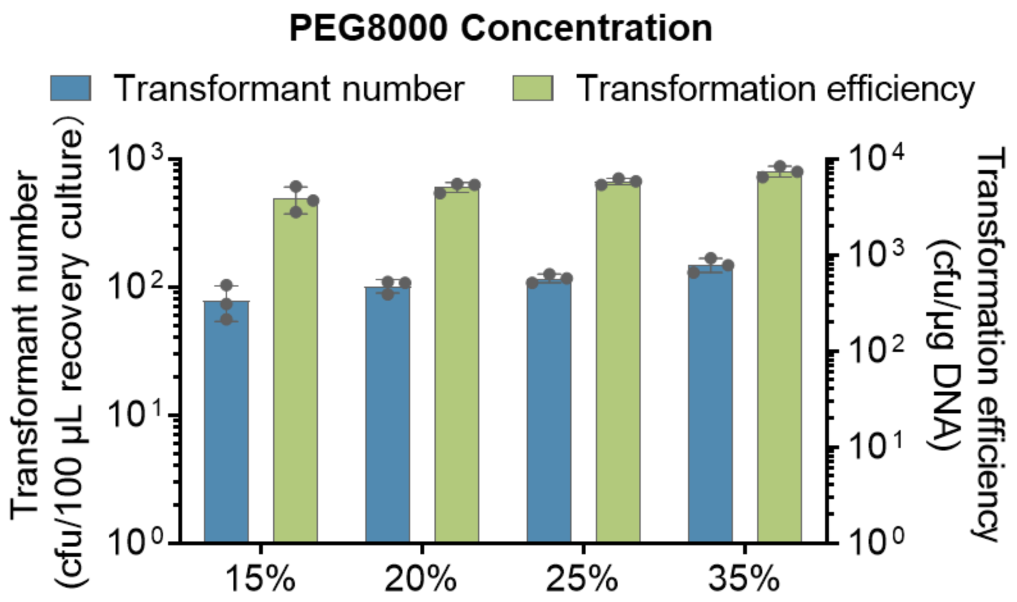


**Supplementary Fig. 2.** Optimization of PEG8000 concentration for electro-transformation. Values and error bars represent the mean ± standard deviation (s.d.) of three biological replicates (n = 3).

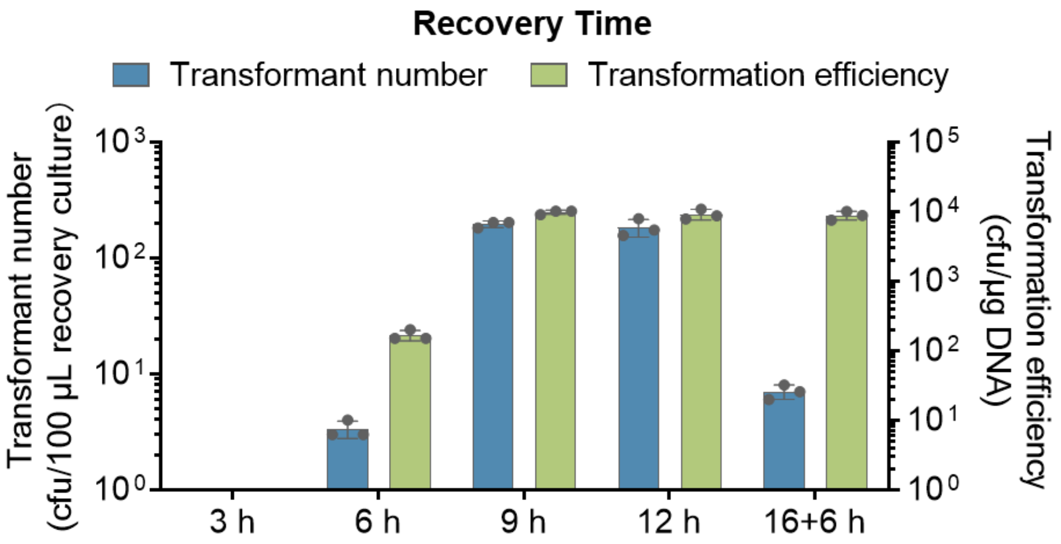


**Supplementary Fig. 3.** Optimization of recovery time for electro-transformation. Values and error bars represent the mean ± standard deviation (s.d.) of three biological replicates (n = 3).

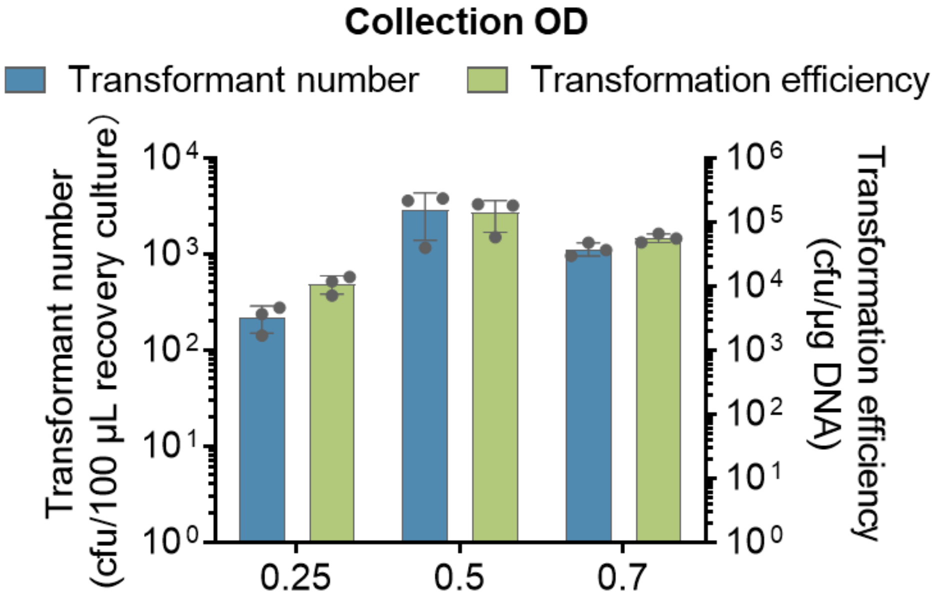


**Supplementary Fig. 4.** Optimization of collection OD for preparing competent cells for electro-transformation. Values and error bars represent the mean ± standard deviation (s.d.) of three biological replicates (n = 3).

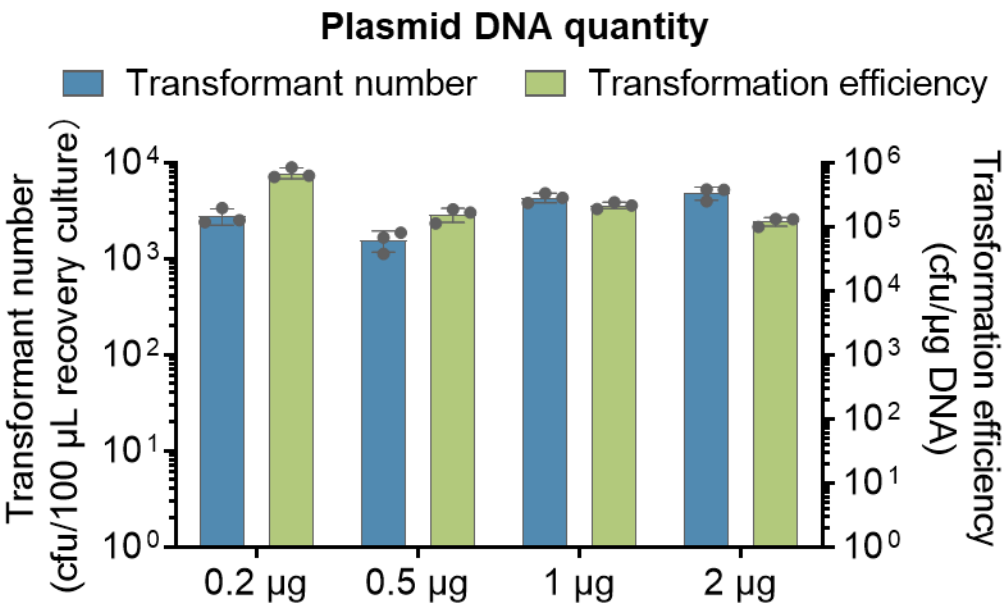


**Supplementary Fig. 5.** Optimization of DNA quantity for electro-transformation. Values and error bars represent the mean ± standard deviation (s.d.) of three biological replicates (n = 3).

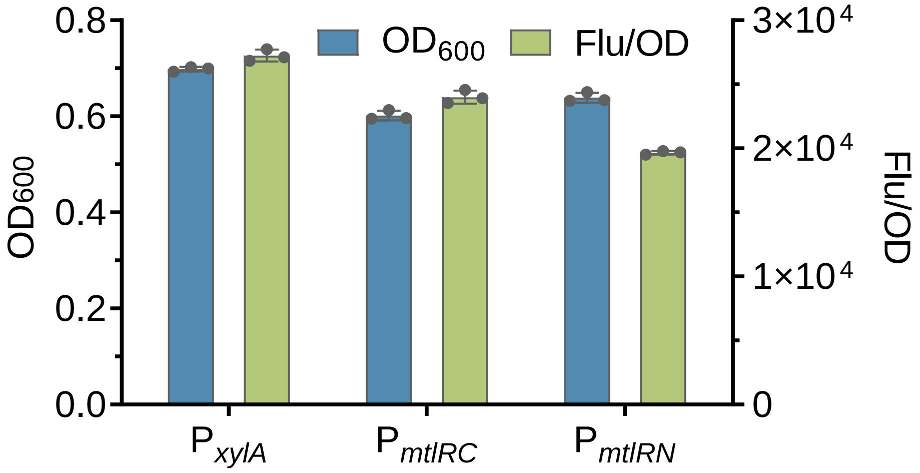


**Supplementary Fig. 6. Basal expression levels of P*xylA* and P*mtlRC* compared to P*mtlRN* in *E. coli*.** Values and error bars represent the mean ± standard deviation (s.d.) of three biological replicates (n = 3).

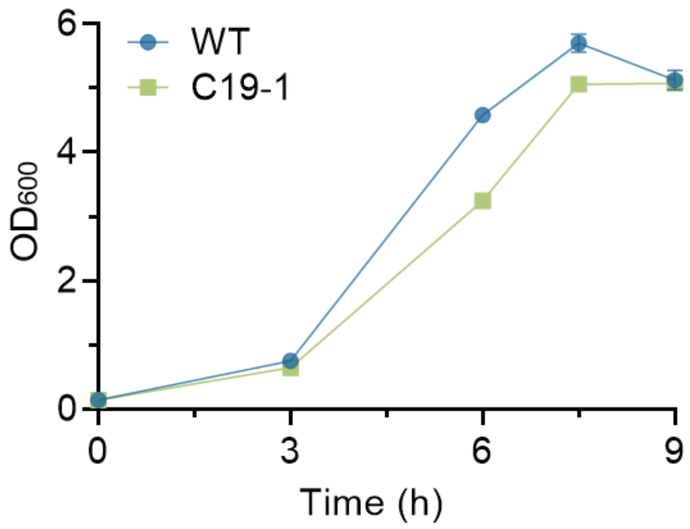


**Supplementary Fig. 7. Cell growth of the wild-type strain and pBM19-integrated strain C19-1 in TSB rich medium.** Values and error bars represent the mean ± standard deviation (s.d.) of three biological replicates (n = 3).

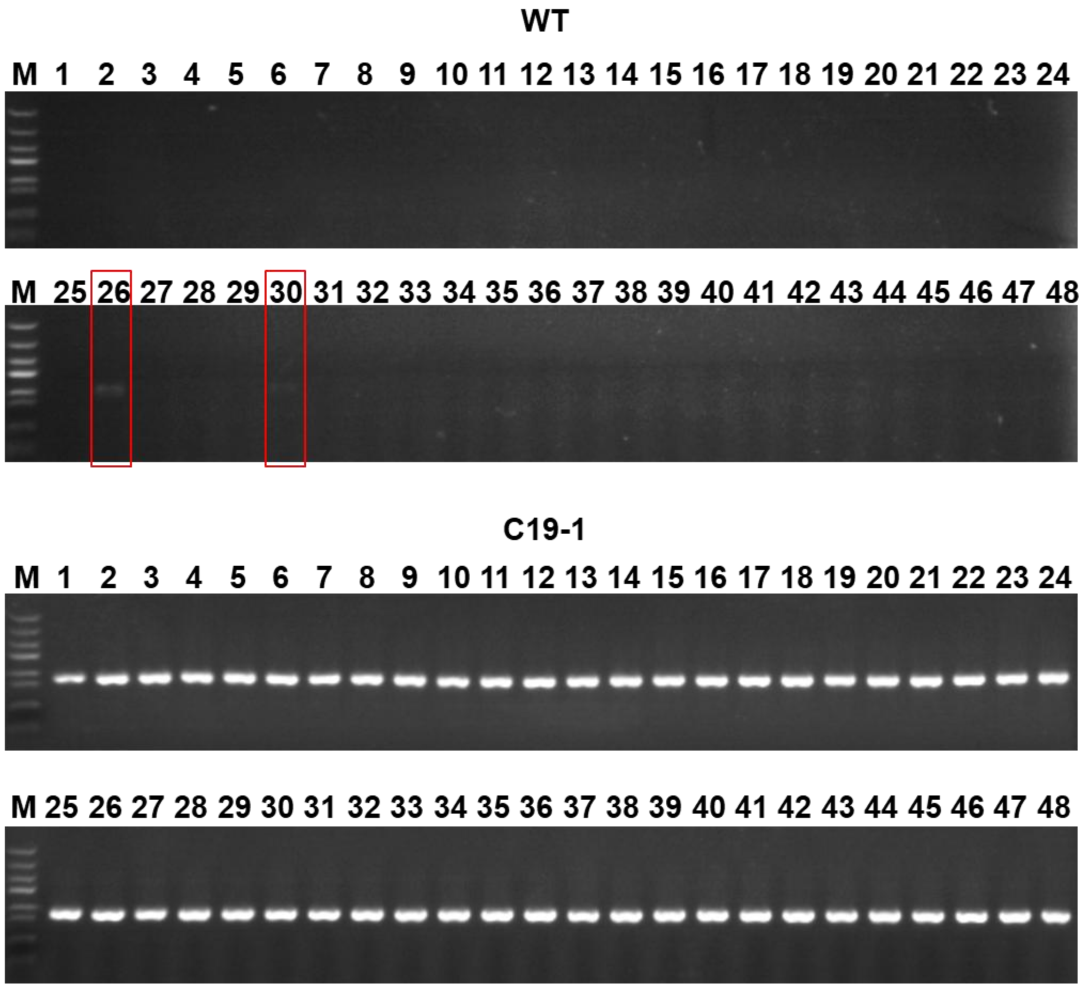


**Supplementary Fig. 8. PCR verification of *rpe*$^{P}$ gene of single colonies isolated from the 10$^{th}$ passage.** The two positive colonies from the culture of the wild-type strain are marked with red boxes. All the tested 48 colonies from the culture of the C19-1 strain are positive.

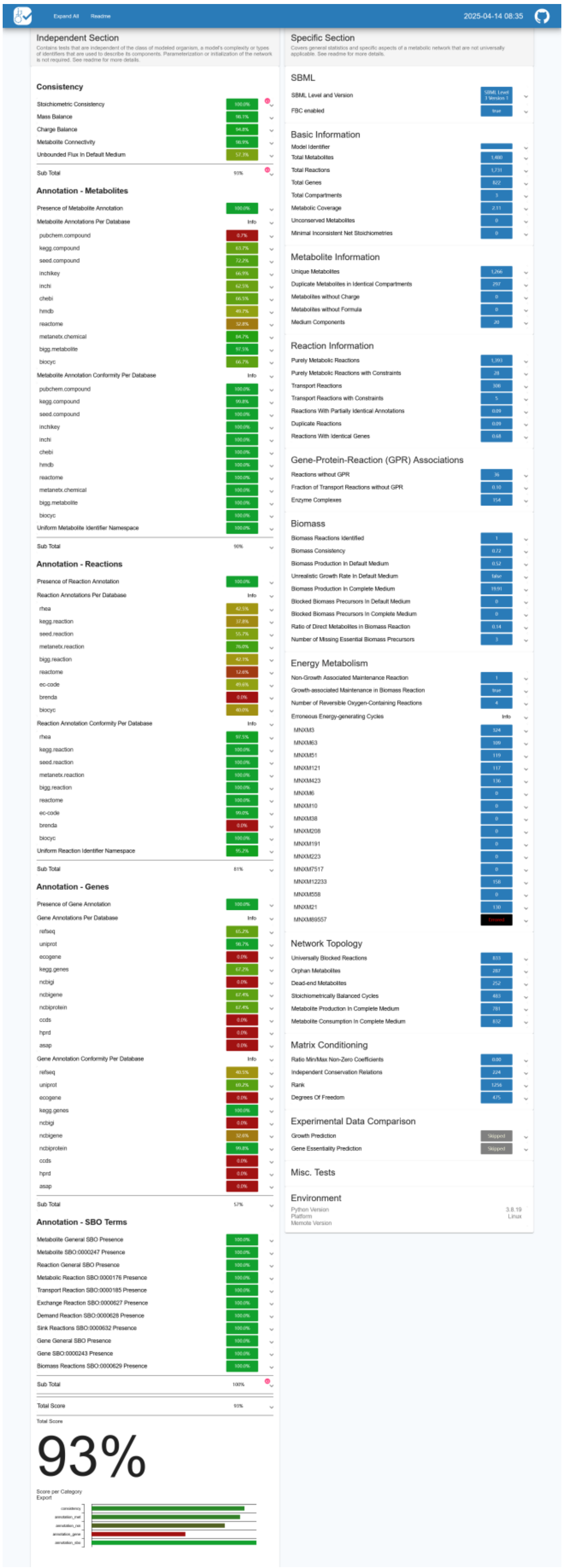

**Supplementary Fig. 9. The MEMOTE comprehensive score of *i*BM822.**

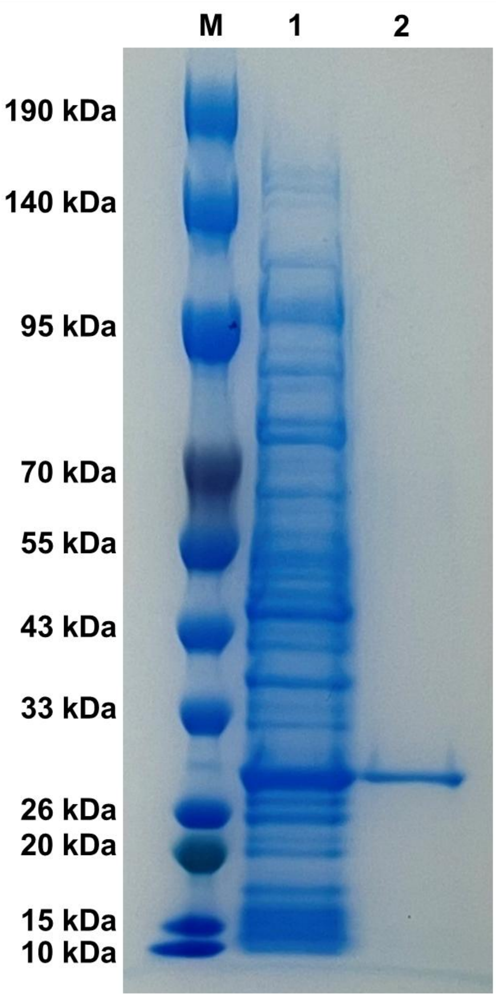


**Supplementary Fig. 10. Purification of recombinant ArgB of *B. methanolicus*.** M, protein marker. 1, supernatant of crude extract of *E. coli* BL21 (DE3) (pET-21a-*argB*). 2, purified ArgB protein.
